# GLI3 Processing by the Primary Cilium Regulates Muscle Stem Cell Entry into G_Alert_

**DOI:** 10.1101/2020.12.07.415273

**Authors:** Caroline E. Brun, Marie-Claude Sincennes, Alexander Y.T. Lin, Derek Hall, William Jarassier, Peter Feige, Morten Ritso, Fabien Le Grand, Michael A. Rudnicki

## Abstract

Satellite cells are required for the growth, maintenance, and regeneration of skeletal muscle. Quiescent satellite cells possess a primary cilium, a structure that regulates the processing of the GLI family of transcription factors. Here we find that GLI3, specifically, plays a critical role in satellite cell activation. Primary cilia-mediated processing of GLI3 is required to maintain satellite cells in a G_0_ dormant state. Strikingly, satellite cells lacking GLI3 enter G_Alert_ in the absence of injury. Furthermore, GLI3 depletion or inhibition of its processing stimulates symmetrical division in satellite cells and expansion of the stem cell pool. As a result, satellite cells lacking GLI3 display rapid cell-cycle entry, increased proliferation and augmented self-renewal, and markedly enhanced long-term regenerative capacity. Therefore, our results reveal an essential role for primary cilia processing of GLI3 in regulating muscle stem cell activation and fate.

## INTRODUCTION

Adult muscle stem cells, or satellite cells, reside within their niche between the basal lamina and myofiber sarcolemma in a reversible G_0_ quiescent state^1^. Upon muscle damage, the niche is modified, inducing the transition from quiescence to activation in satellite cells. This process triggers mechano-property changes, migration, metabolic activation, increased RNA transcription and protein synthesis and cell cycle entry^2-6^. Once activated, satellite cells proliferate extensively to generate myogenic progenitors that differentiate and fuse to repair the injured myofibers, while a subset of satellite cells self-renew and return to quiescence to replenish the stem cell pool^7,8^. Hence, the ability to self-renew and reversibly enter quiescence is a hallmark of satellite cells that ensures proper muscle repair throughout life. Interestingly, in response to extrinsic cues, satellite cells can dynamically transit from a G_0_ to a G_Alert_ quiescent state, which is mediated by mTORC1 signaling^6,9-11^. Whether other pathways regulate this poised activation state that confers enhanced regenerative capacity to the G_Alert_ satellite cells remains unestablished.

A high proportion of quiescent satellite cells harbor a primary cilium, which rapidly disassembles upon activation and reassembles preferentially in self-renewing satellite cells^12^. The primary cilium is a small, non-motile, microtubule-based structure anchored to a cytoplasmic basal body that protrudes from cells in G_0_^13^. It acts as a nexus for cellular signaling, most notably Hedgehog signaling^14,15^, wherein cilia-mediated processing of the GLI-family of transcription factors is required for Hedgehog signal transduction^16-18^. Although there are three GLI transcription factors (GLI1-3), only GLI3 contains a potent N-terminal repressor domain.

In the absence of Hedgehog ligands, GLI3 is sequentially phosphorylated at the base of the cilium, first by the cAMP-dependent protein kinase A (PKA) and then by glycogen synthase kinase 3 (GSK3) and casein kinase 1 (CK1)^19-23^. GLI3 phosphorylation promotes its proteolytic cleavage, converting it to a repressor form. Accordingly, decreased PKA activity leads to the activation of Hedgehog signaling independently of Hedgehog ligand-receptor binding^24-26^. Hedgehog ligand-receptor binding induces the accumulation of GLI3 at the cilium tip, thereby limiting its phosphorylation and cleavage^16,21^. Thus, the primary cilium controls the balance between the full-length activator (GLI3FL) and cleaved repressor (GLI3R), which typically dictates the activation state of Hedgehog signaling.

Although studies have demonstrated that primary cilia act as a signaling hub, their role in satellite cell function remains unknown. Here, we identify GLI3 as a mediator of the cell-autonomous, cilia-related control of satellite cell activation. Using a conditional knockout strategy, we show that ciliary processing of GLI3 to the repressor form (GLI3R) controls the G_0_ to G_Alert_ transition of quiescent muscle stem cells through regulation of mTORC1 signaling, independent of canonical Hedgehog downstream target gene expression. Moreover, we find that GLI3R regulates satellite cell commitment by promoting asymmetric cell division and that genetic ablation of *Gli3* promotes symmetric cell expansion and improves muscle repair. We conclude that primary cilia-mediated GLI3 processing controls the activation and regenerative capacity of muscle stem cells. Furthermore, we identify GLI3 as a potential therapeutic target to promote muscle stem cell engraftment in conditions such as Duchenne muscular dystrophy.

## RESULTS

### GLI3 is the major GLI-family transcription factor expressed in muscle stem cells

Satellite cells harbor a primary cilium^12,27,28^, yet the cilia-mediated pathways regulating satellite cell function remain uncharacterized. As the Hedgehog pathway requires the primary cilium for its transduction^14^, we hypothesized that cilia-mediated Hedgehog signaling regulates satellite cell function.

We first performed RNA-sequencing on freshly isolated satellite cells from resting and injured muscles. Although FACS method induces partial activation^29-31^, we refer to the freshly isolated satellite cells as quiescent satellite cells (QSCs) in the manuscript. In addition, the mixed population of proliferating satellite cells and progenitors isolated from 3 days post-cardiotoxin (CTX)-injured muscles will be referred as to activated satellite cells (ASCs)^32^. We observed that the components of canonical Hedgehog signaling, namely the receptor *Patched1 (Ptch1)*, the signal transducer *Smoothened (Smo)* and the three transcriptional effectors *Gli1*, *Gli2* and *Gli3*, are expressed in quiescent satellite cells (**Supplementary Fig. 1a**). However, expression of conserved GLI-target genes^33^, such as *Gli1*, *Ptch1, Ptch2, Hhip* and *Bcl2*, is significantly downregulated as satellite cells transit from quiescence to activation (**Supplementary Data file 1; Supplementary Fig. 1b**). The increased expression of *Cdk6, Ccnd2* and *Ccnd1* is likely due to the cycling state of the ASCs. Together, these results indicate that canonical Hedgehog signaling is turned off as QSCs become activated. Of note, the transcriptional effectors *Gli1* and *Gli2* are strikingly downregulated, while *Gli3* expression is significantly increased in ASCs (**Supplementary Data file 1; Supplementary Fig. 1b**). RT-qPCR on QSCs, ASCs, proliferating myoblasts and differentiated myotubes further confirmed that only *Gli3* is enriched in ASCs and proliferating myoblasts (**Supplementary Fig. 1c**).

**Figure 1.**
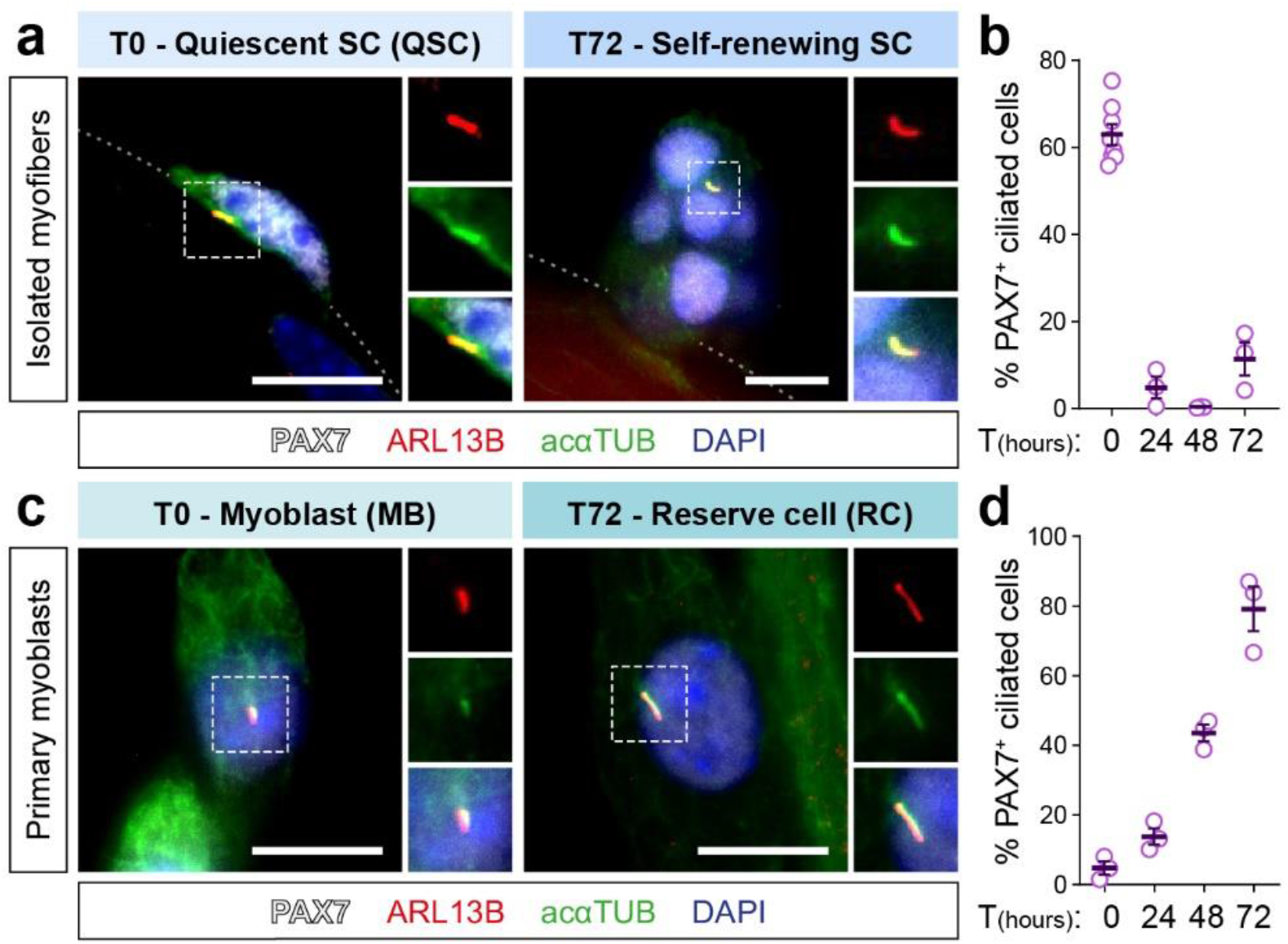
Primary cilia are dynamically regulated in muscle cells. **a**) Representative immunofluorescence staining of PAX7 (white), ARL13B (red), acetylated alpha-TUBULIN (acαTUB, green) and nuclei (blue) showing primary cilia on a quiescent satellite cell (SC) from freshly isolated myofiber (T0) and a self-renewing SC from 72h-cultured myofiber (T72). **b**) Proportion of PAX7^+^ satellite cells harboring a primary cilium on freshly isolated myofibers (QSCs, n = 6) and 24h, 48h and 72h-cultured myofibers (n = 3). **c**) Representative immunostaining of PAX7 (white), ARL13B (red), acαTUB (green) and nuclei (blue) showing primary cilia on a myoblast (MB) (T0) and a reserve cell (RC) (T72). **d**) Proportion of PAX7^+^ primary myoblasts harboring a primary cilium during 72h-differentiation time course (n = 3). Scale bars, 10μm; Error bars, SEM.

### Primary cilia dynamically regulate GLI3 proteolytic processing in myogenic cells

As GLI3 activity relies on the primary cilium^17,21^, we investigated the kinetics of primary cilia assembly and disassembly on cultured myofibers and primary myoblasts. Primary cilia were labelled by immunostaining using antibodies directed against the primary cilium-specific protein ARL13B and acetylated α-TUBULIN (acαTUB), and basal bodies were stained with γ-TUBULIN (γTUB) (**Fig. 1; Supplementary Fig. 1d, e**). In line with previous observations^12^, we found that more than 60% of the satellite cells on freshly myofibers isolated from *extensor digitorum longus*(EDL) muscle have a primary cilium that disassembles during proliferation and reassembles specifically in the PAX7^+^ self-renewing cells (**Fig. 1b**). As well, proliferating myoblasts are rarely ciliated and only the PAX7^+^ ‘reserve’ cells^34^ reassemble a primary cilium upon differentiation (**Fig. 1d**).

Full-length and non-PKA phosphorylated GLI3 (GLI3FL) localizes in the primary cilium, while the cleaved repressor GLI3R is found at the basal body of the cilium where its proteolytic processing occurs^16,21,35,36^. To confirm that similar regulation occurs in myogenic cells, we treated primary myoblasts with SAG, a SMOOTHENED agonist that induces accumulation of GLI3FL in the primary cilium, thus abrogating its proteolytic processing^21,36^ (**Supplementary Fig. 2a-g**). Consequently, SAG-induced Hedgehog stimulation decreases the level of GLI3R, leading to the upregulation of the two well-known GLI-target genes, *Gli1* and *Ptch1* (**Supplementary Fig. 2a-f**). To assess the effects of negative regulation, myoblasts were treated with forskolin (FSK), an adenylyl cyclase activator that stimulates PKA^21,35^. FSK treatment prevents GLI3 ciliary accumulation, promotes GLI3 proteolytic cleavage and *Gli1* and *Ptch1* expression is decreased (**Supplementary Fig. 2a-e, g**). Therefore, we conclude that cilia and PKA-mediated regulation of GLI3 processing is conserved in myogenic cells.

We then assessed the temporal coordination of muscle cell ciliation and both GLI3 localization and proteolytic processing by immunostaining and Western blot. In ciliated QSCs on freshly isolated myofibers (T0), GLI3 antibody clearly labels the basal body, where PKA is expressed (**Fig. 2a**), suggesting that GLI3 is expressed as a repressor. As satellite cells activate and lose primary cilia, GLI3 localizes in the cytoplasm and around the microtubule-organizing centers (**Fig. 2a**). In cultured primary cells, although primary ciliation is rare due to their proliferative activity, 65% of ciliated myoblasts display GLI3 accumulation at the axoneme and the tip of primary cilia (**Fig. 2b, c**). In non-ciliated myoblasts, GLI3 is mainly found in the cytoplasm, as observed in ASCs on myofibers (**Fig. 2a, b**). GLI3FL protein levels are the highest in proliferating myoblasts and decrease during myogenic differentiation concomitantly with the ratio of GLI3FL/GLI3R (**Fig. 2d-g**), which correlates with the kinetics of ciliation and ARL13B expression (**Fig. 2b-g**). Finally, as myoblasts undergo differentiation, GLI3 progressively transits from the axoneme to the basal body of primary cilia and at 72h, most of the PAX7^+^‘reserve’ cells have a cilium and maintain high GLI3R, whereas myotubes have no primary cilium and do not express GLI3 (**Fig. 2**).

**Figure 2.**
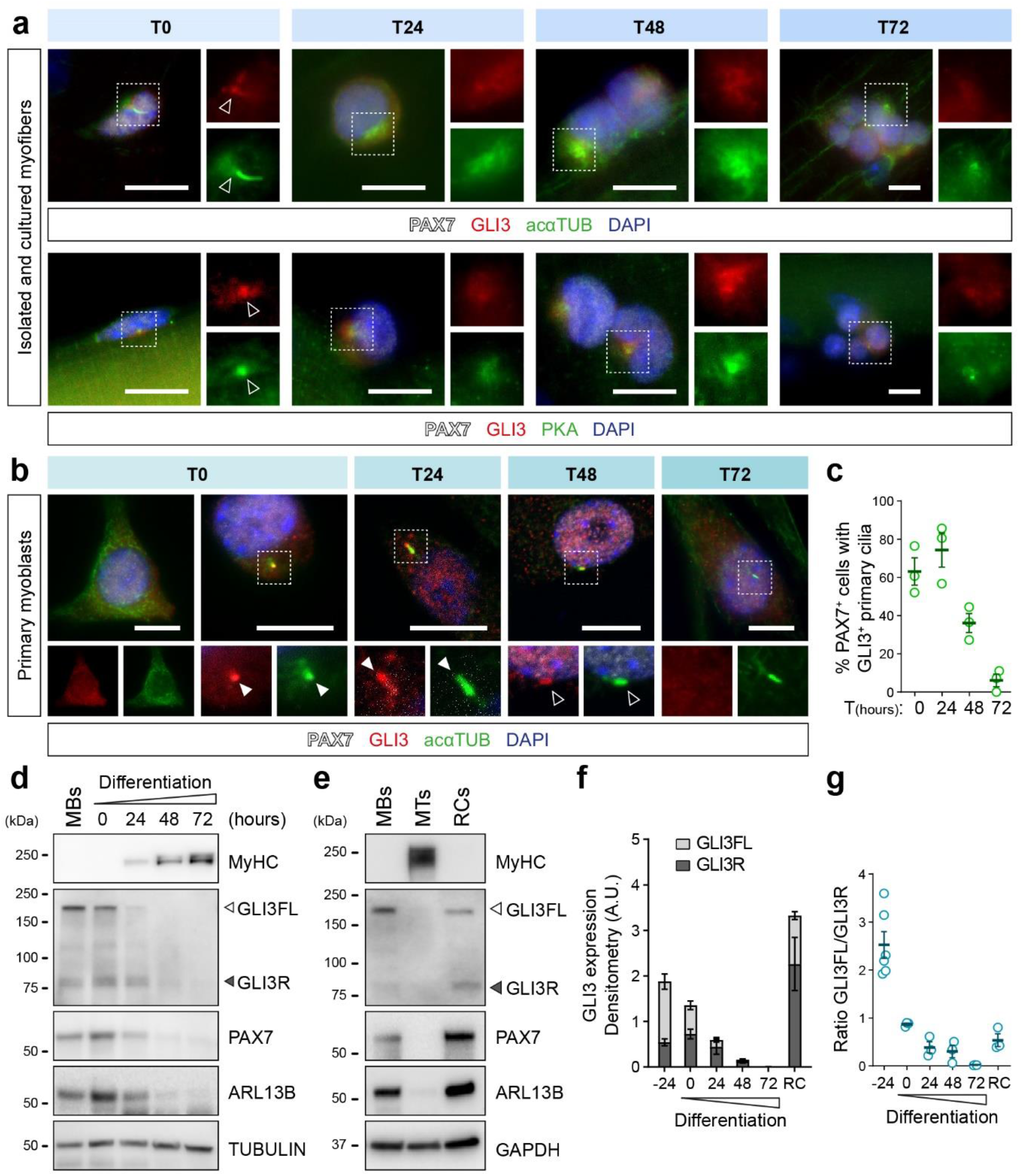
GLI3 subcellular localization and proteolytic processing rely on primary cilia dynamics. **a**) Immunostaining showing GLI3 (red) localization in PAX7^+^ (white) quiescent (T0), activated (T24) and proliferating (T48-72) satellite cells on isolated and cultured myofibers. Upper panel, Acetylated α-TUBULIN (acαTUB, green) stains the primary cilium (T0) and the microtubule-organizing centers (T24-72). Lower panel, PKA (green) localizes at and around the centrosome areas. **b**) Immunostaining of GLI3 (red) in primary cilia labeled by acetylated α-TUBULIN (acαTUB, green) of PAX7^+^ primary myoblasts (white) during their differentiation time course. **c**) Proportion of PAX7^+^ cells that display a GLI3^+^ primary cilium during myoblast differentiation time course (n= 3). **d**) Immunoblot analysis of the ciliary membrane marker ARL13B, GLI3 full-length (GLI3FL) and repressor (GLI3R), in proliferating myoblasts and from 0h to 72h of differentiation. PAX7 and MyHC (myosin heavy chain) are used to monitor myogenic differentiation. TUBULIN is used as a loading control. **e**) Immunoblot analysis of ARL13B, GLI3FL and GLI3R in myoblasts (MBs), 72h-differentiated myoblasts or myotubes (MTs) and reserve cells (RCs). Both MBs and RCs express PAX7 while MTs express the myogenic differentiation marker MyHC. GAPDH is used as a loading control. **f**) Densitometric analysis of the level of GLI3FL and GLI3R relative to TUBULIN signals of 6 (−24h) and 3 biological replicates (0h-72h). **g**) Ratio of GLI3FL/GLI3R relative to GAPDH (−24h, n = 6; 0h-72h, n = 3). White arrows indicate the tip of the primary cilium; Empty arrows indicate the basal body; Scale bar, 10μm; Error bars, SEM.

In the absence of ligand stimulation, primary cilia trigger GLI3 proteolytic processing, restraining the Hedgehog pathway in an off state^13^. To assess whether primary cilia-mediated processing of GLI3 controls the output of Hedgehog signaling in myogenic cells, primary myoblasts were treated with either siRNA targeting *Gli3*, or with siRNA against *Ift88* to disrupt primary cilia assembly and consequently GLI3 processing^18,37^ (**Supplementary Fig. 2h-l**). Knocking-down *Ift88* decreases GLI3R levels, consequently increasing GLI3R/GLI3FL ratio and *Gli1* and *Ptch1* expression (**Supplementary Fig. 2h-k**). Interestingly, *Gli3* siRNA treatment induces similar upregulation of the two GLI-target genes (**Supplementary Fig. 2l**), indicating that GLI3FL is dispensable for GLI-target gene expression and that the predominant regulatory form is the repressor.

These results collectively show that GLI3 is processed to a repressor in ciliated QSCs. Transient primary cilia disassembly during cell cycle entry and proliferation abolishes GLI3R processing, promoting ciliary GLI3FL accumulation. Upon differentiation, myoblasts reassemble a primary cilium that will be only maintained in the self-renewing ‘reserve’ cell population, which then expresses GLI3R at the base of the cilium. Thus, our results demonstrate that the ratio of GLI3FL/GLI3R relies on cilia dynamics and is regulated independently of Hedgehog exogenous signals as muscle stem cells progress through the myogenic lineage.

### Loss of GLI3R does not affect Hedgehog signaling in satellite cells

To better characterize the role of GLI3R in regulating satellite cell function, we generated *Pax7^CE/+^; Gli3^+/+^; R26R^YFP^* and *Pax7^CE/+^; Gli3^fl/fl^ R26R^YFP^* (hereafter referred to as *Gli3*^+/+^ and *Gli3*^Δ/Δ^ mice, respectively), in which *Gli3* was excised in satellite cells that are simultaneously labelled by YFP upon tamoxifen induction (**Supplementary Fig. 3a**). The efficiency of *Gli3* excision was confirmed in QSCs and their myogenic descendants (ASCs, MBs and MTs) (**Supplementary Fig. 3b, c**).

Uninjured resting muscle exhibit no gross histological abnormalities upon tamoxifen treatment (**Fig. 3a**). However, *Gli3*^Δ/Δ^ muscles were found to have a transient increase in the number of satellite cells following tamoxifen injections (**Fig. 3b**). This observation led us to hypothesize that *Gli3* deletion in satellite cells could be inducing a break from quiescence. During homeostasis, satellite cells reside in their niche, which maintains their quiescence, but then migrate outside the basal lamina upon activation^2,5^. Analyzing muscle cross-sections and myofibers post-tamoxifen injection showed that both *Gli3*^+/+^ and *Gli3*^Δ/Δ^ satellite cells are located in their niche, underneath the basal lamina (**Supplementary Fig. 3d-f**) and that *Gli3*^+/+^ and *Gli3*^Δ/Δ^ mice exhibit similar proportion of ciliated cells (**Fig. 3c, d**). Together, these results indicate that *Gli3*^Δ/Δ^ satellite cells are inherently in quiescent state and suggests that Cre-mediated *Gli3* deletion transiently activates satellite cells, increasing their number over weeks after tamoxifen treatment.

**Figure 3.**
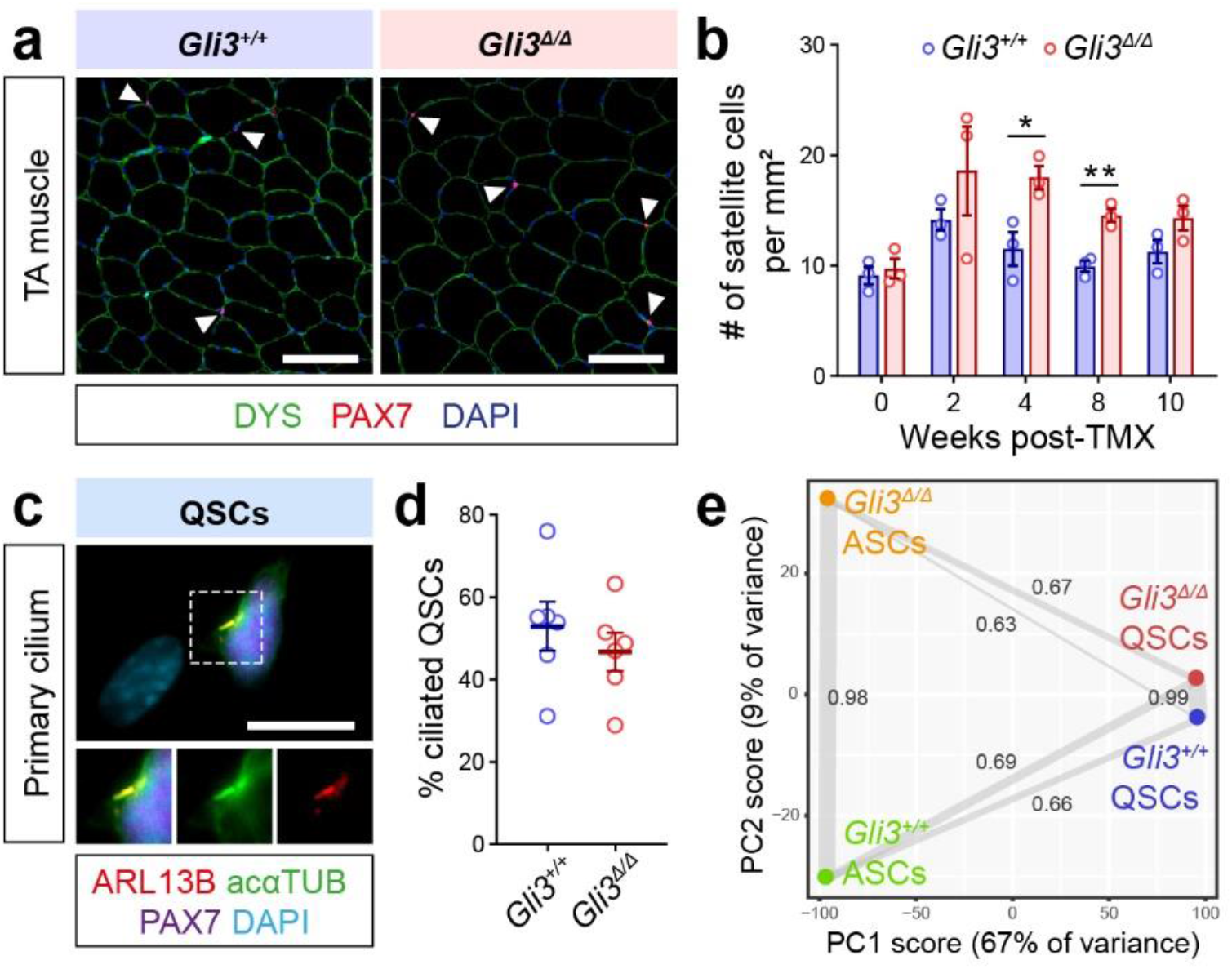
GLI3 controls the quiescence of satellite cells. **a**) Representative immunofluorescence picture showing transversal cross-sections of resting TA muscle in *Gli3*^+/+^ and *Gli3*^Δ/Δ^ mice. PAX7 (red) labels the satellite cells and DYS (green) delineates the myofibers. Nuclei are stained with DAPI (blue). Scale bars represent 50μm. **b**) Quantification of the number of PAX7^+^ satellite cells per mm^2^ in resting TA muscle from *Gli3*^+/+^ and *Gli3*^Δ/Δ^ mice following tamoxifen treatment (n = 3 males). **c**) Primary cilia are immunostained with ARL13B (red) and acαTUB (green) and satellite cells with PAX7 (purple). Nuclei are labeled with DAPI (blue). Scale bars represent 10μm. **d**) Proportions of ciliated satellite cells on freshly isolated EDL myofibers from *Gli3*^+/+^ and *Gli3*^Δ/Δ^ mice (n = 6, 3 males and 3 females for each genotype). **e**) Principal component analysis (PCA) of global transcriptomes of *Gli3*^+/+^ and *Gli3*^Δ/Δ^ quiescent (QSCs) and activated satellite cells (ASCs) and Pearson’s values showing the correlation between samples. Each dot represents the mean of 3 biological samples. Error bars, SEM; \**p* < 0.05; \*\*\**p* < 0.001

To further characterize the consequences of *Gli3* deletion, we performed RNA-sequencing on *Gli3*^+/+^ and *Gli3*^Δ/Δ^ QSCs and ASCs and compared their transcriptome (**Fig. 3e; Supplementary Fig. 3g**). Principle component analysis (PCA) of the transcriptional profiles of *Gli3*^+/+^ and *Gli3*^Δ/Δ^ SCs revealed two distinct groups based on the first component axis (PC1), distinguishing the QSCs from the ASCs (**Fig. 3e**). PCA also revealed a high correlation between *Gli3*^+/+^ and *Gli3*^Δ/Δ^ QSCs, corroborating the idea that the *Gli3*^Δ/Δ^ SCs are predominantly quiescent. Interestingly, the correlation between the QSC and ASC transcriptional signatures was slightly increased in the *Gli3*^Δ/Δ^ background (0.67) than the *Gli3*^+/+^ (0.66) (**Fig. 3e**), suggesting that the *Gli3*^Δ/Δ^ QSCs may actually exhibit some features of satellite cell activation.

Surprisingly, *in silico* analysis did not highlight any enrichment for canonical Hedgehog signaling (**Fig. 4; Supplementary Fig. 4a-d**). RT-qPCR further confirmed that the GLI-target genes, *Gli1* and *Ptch1*, have the same level of expression in *Gli3*^+/+^ and *Gli3*^Δ/Δ^ QSCs and ASCs (**Supplementary Fig. 4e, f**), suggesting that GLI3 does not repress its canonical target genes in satellite cells. Instead, Gene Ontology (GO) analysis of the 41 upregulated transcripts in *Gli3*^Δ/Δ^QSCs revealed an enrichment for biological process terms related to glucose and insulin signaling (**Fig. 4a; Supplementary Data file 2**). In *Gli3*^Δ/Δ^ ASCs, 144 transcripts are significantly upregulated and correlate with GO terms associated with regulation of G1/S phase transition and cell cycle (**Fig. 4b; Supplementary Data file 2**). The gene set enrichment analysis (GSEA) also confirmed the activation of cell cycle-related pathways (MYC/E2F targets, G2M checkpoint, DNA repair) in ASCs (**Fig. 4d**). Interestingly, mTORC1 signaling was identified among the active pathways in both *Gli3*^Δ/Δ^ QSCs and ASCs (**Fig. 4c, d**). mTORC1 signaling activity drives the G_0_- to-G_Alert_ transition in quiescence^6^ and is required for satellite cell proliferation and fusion upon activation^38^. Therefore, we hypothesized that *Gli3*^Δ/Δ^ satellite cells have transitioned into G_Alert_.

**Figure 4.**
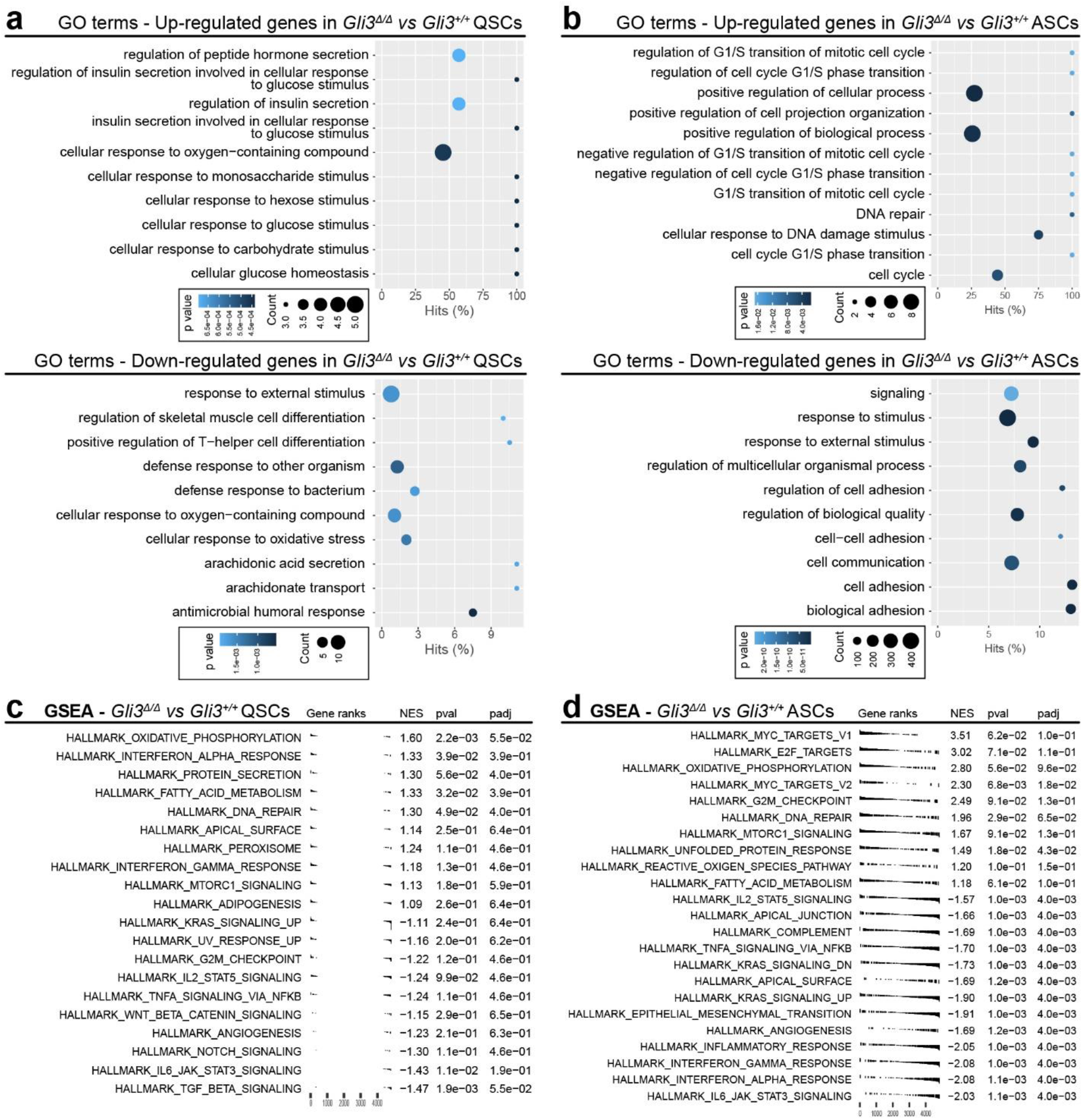
Gli3 deletion in satellite cells activates mTORC1 signaling. **a**) GO term analysis Gene Ontology (GO) term enrichment (Biological process) for the up-regulated and down-regulated genes in *Gli3*^Δ/Δ^ QSCs compared to *Gli3*^+/+^ QSCs. **b**) GO term analysis Gene Ontology (GO) term enrichment (Biological process) for the up-regulated and down-regulated genes in *Gli3*^Δ/Δ^ ASCs compared to *Gli3*^+/+^ ASCs. **c**) GSEA of *Gli3* QSCs compared to *Gli3*^+/+^ QSCs. **d**) GSEA of *Gli3* ASCs compared to *Gli3*^+/+^ ASCs.

### GLI3R regulates the entry of quiescent satellite cells into G_Alert_

In contrast to quiescent G_0_ satellite cells, G_Alert_ satellite cells display an increase in cell size, transcriptional activity and mitochondrial metabolism, and are poised to activate faster in response to injury^6^. Imaging flow cytometry revealed that *Gli3*^Δ/Δ^ QSCs display an increase in size compared to *Gli3*^+/+^ QSCs (**Fig. 5a-c; Supplementary Fig. 5a**). *Gli3*^Δ/Δ^ QSCs have increased PyroninY staining, consistent with increased transcriptional activity, and higher mitochondrial mass, in line with our RNA-sequencing data showing an enrichment in genes related to oxidative phosphorylation (**Fig. 5d-g; Supplementary Fig. 5b**). None of these features, however, overlaps with the profiles observed in ASCs, confirming that the *Gli3*^Δ/Δ^ QSCs are not fully activated (**Fig. 5b-g**).

**Figure 5.**
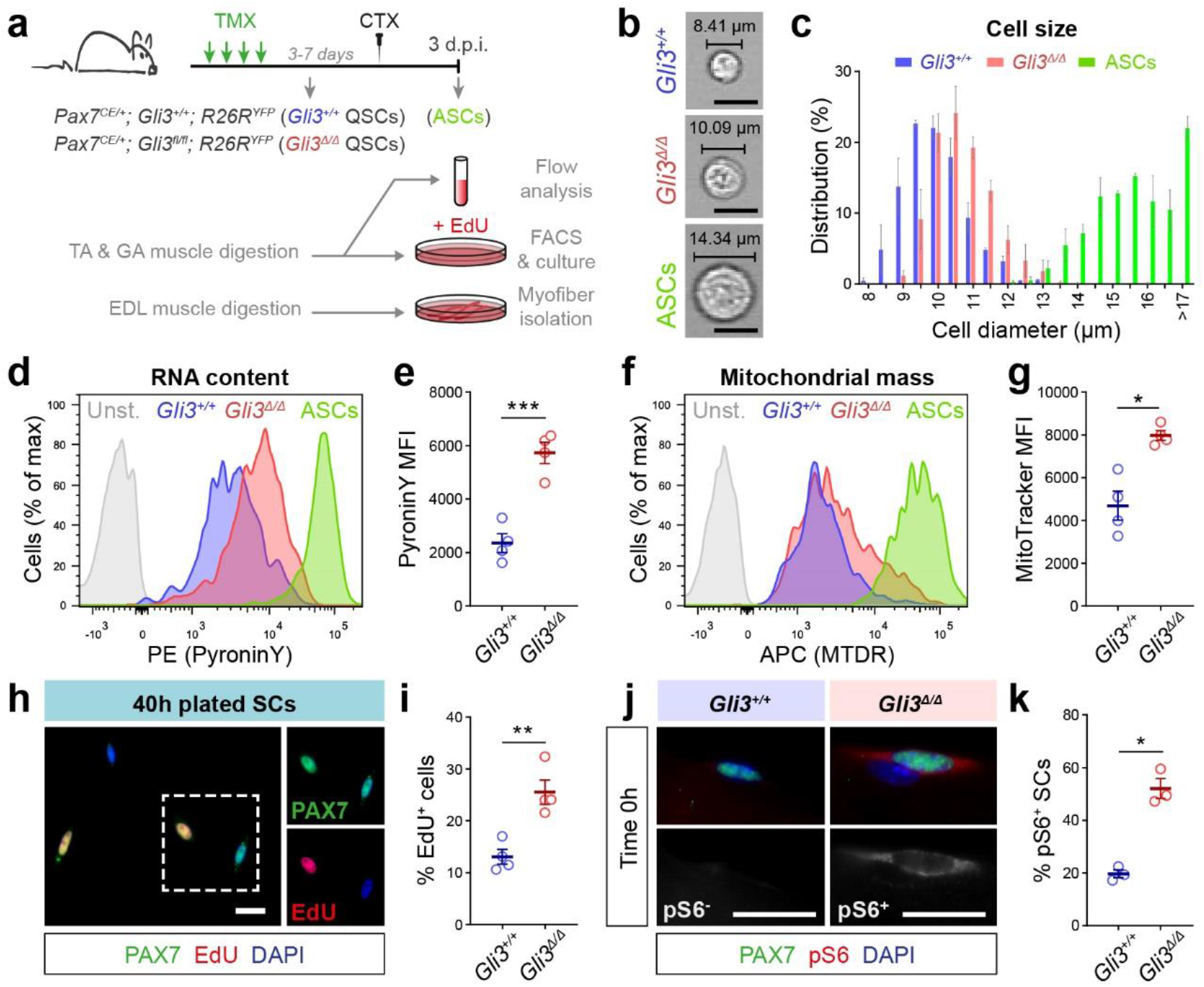
Satellite cells in Gli3^Δ/Δ^ uninjured muscle are in G_Alert_. **a**) Experimental design. Satellite cells (SCs) from TA and *gastrocnemius* (GA) muscles were analyzed for cell size, RNA content and mitochondrial mass. FACS-isolated SCs were plated 40h with EdU to analyze cell cycle entry. Phosphorylation status of S6 ribosomal protein was analyzed on freshly isolated EDL myofibers. **b**) Representative images of freshly sorted *Gli3*^+/+^ and *Gli3*^Δ/Δ^ quiescent satellite cells (QSCs) and *Gli3*^+/+^ activated satellite cells (ASCs). **c**) Distribution of *Gli3*^+/+^ QSCs, *Gli3*^Δ/Δ^ QSCs and *Gli3*^+/+^ ASCs according to their diameter (n = 3 males, >100 cells per mouse). **d**) Representative flow cytometry plot of *Gli3*^+/+^QSCs, *Gli3*^Δ/Δ^ QSCs and *Gli3*^+/+^ ASCs stained for PyroninY (Unst. = unstained). **e**) Mean fluorescence intensity (MFI) of PyroninY staining in *Gli3*^+/+^ and *Gli3*^Δ/Δ^ QSCs (n = 4 males). **f**) Representative flow cytometry plot of *Gli3*^+/+^ QSCs, *Gli3*^Δ/Δ^ QSCs and *Gli3*^+/+^ ASCs stained for MitoTracker Deep Red (MTDR). **g**) MFI of MitoTracker staining in *Gli3*^+/+^ and *Gli3*^Δ/Δ^ QSCs (n = 4 males). **h**) Immunostaining of SCs with PAX7 (green) and DAPI (blue) showing EdU (red) incorporation 40h post-isolation. **i**) Proportion of EdU^+^ SCs in *Gli3*^+/+^ and *Gli3*^Δ/Δ^ mice (n = 4, 3 males and 1 female). **j**) Immunofluorescence of phospho-S6 (pS6, red) in SCs (PAX7, green) on freshly isolated myofibers from *Gli3*^+/+^ and *Gli3*^Δ/Δ^ mice. **k**) Proportion of pS6^+^ SCs in *Gli3*^+/+^ and *Gli3*^Δ/Δ^ mice (n = 3 females). Scale bars, 10μm; Error bars, SEM; \**p* < 0.05; \*\**p* < 0.01; \*\*\**p* < 0.001.

To assess their ability to enter the cell cycle, *Gli3*^+/+^ and *Gli3*^Δ/Δ^ QSCs were cultured immediately after FACS-isolation in the presence of EdU for 40h (**Fig. 5h**). The proportion of EdU^+^ cells was significantly higher in the *Gli3*^Δ/Δ^ culture (**Fig. 5i**), demonstrating that *Gli3*^Δ/Δ^ QSCs enter the cell cycle faster. This was further confirmed using freshly isolated myofibers cultured with EdU for 24h, where *Gli3*^Δ/Δ^ myofibers show a higher proportion of EdU^+^ SCs compared to *Gli3*^+/+^(**Supplementary Fig. 5c-e**).

As mTORC1 signaling regulates the G_0_-to-G_Alert_ transition in muscle stem cells^6^, we analyzed the phosphorylation of the ribosomal protein S6 as a marker for mTORC1 activation in freshly isolated myofibers from *Gli3*^+/+^ and *Gli3*^Δ/Δ^ mice (**Fig. 5j**). Remarkably, we observed that more than 50% of the *Gli3*^Δ/Δ^ SCs display staining for phospho-S6, while only 20% of *Gli3*^+/+^ SCs are phospho-S6^+^ (**Fig. 5k**). Altogether, these findings demonstrate that GLI3R represses mTORC1 signaling to maintain satellite cells in G_0_ quiescence and that loss of GLI3R induces a transition to G_Alert_.

It has previously been demonstrated that G_0_ and G_Alert_ satellite cells share similar engraftment efficiency and capacity for self-renewal^6^. To confirm that *Gli3*^Δ/Δ^ satellite cells also retain stemness properties, we performed transplantation experiments of *Gli3*^+/+^ and *Gli3*^Δ/Δ^ QSCs into irradiated TA muscles of immunocompromised MDX mice, a mouse model for Duchenne muscular dystrophy (**Supplementary Fig. 5f**). Remarkably, *Gli3*^Δ/Δ^ SCs exhibit enhanced capacity to engraft and self-renew as they generate 3.5 times more DYS^+^/YFP^+^ myofibers and PAX7^+/^YFP^+^ satellite cells than the *Gli3*^+/+^ SCs (**Supplementary Fig. 5g-j**). Given that enhanced engraftment is not a phenotype associated with G_Alert_ satellite cells, this indicates that *Gli3* deletion must affect other aspects of satellite cell biology beyond cell cycle regulation and, specifically, may affect satellite cell self-renewal.

### GLI3R regulates the self-renewal of satellite cells

To evaluate the role of GLI3 in regulating satellite cell self-renewal, we utilized the *Myf5^cre^;R26R^YFP^* mouse model. These mice allow for the tracking of the expression of the myogenic determination factor *Myf5*, following the first division of YFP^-^ stem cells that have never expressed *Myf5^7^.* Therefore, they allow analysis of asymmetric divisions and the early commitment of satellite cells^7,39,40^.

To assess the effects of GLI3 depletion or loss of GLI3 processing, we treated single EDL *Myf5^cre^;R26R^YFP^* myofibers with either *Gli3* or *Ift88* siRNA, respectively (**Fig. 6a**). We examined satellite cells immediately after their first cell division at 42h of culture, when the number of asymmetric and symmetric cell doublets reflects either the commitment or the expansion of the stem cell pool, respectively (**Fig. 6b**). Both *Ift88* and *Gli3* siRNA significantly decrease the number of asymmetric cell doublets (**Fig. 6c, g**). Although *Gli3* siRNA does not significantly change the number of symmetric divisions of YFP^-^ satellite cells (**Fig. 6d, h**), both siRNA treatments increase the proportion of YFP^-^ symmetric divisions (**Fig. 6e, i**), without changing the total number of satellite cells per fiber (**Fig. 6f, j**). These data suggest that loss of GLI3 or disruption of its processing through ciliogenesis impairment drive satellite cell first division towards self-renewal and expansion rather than myogenic commitment.

**Figure 6.**
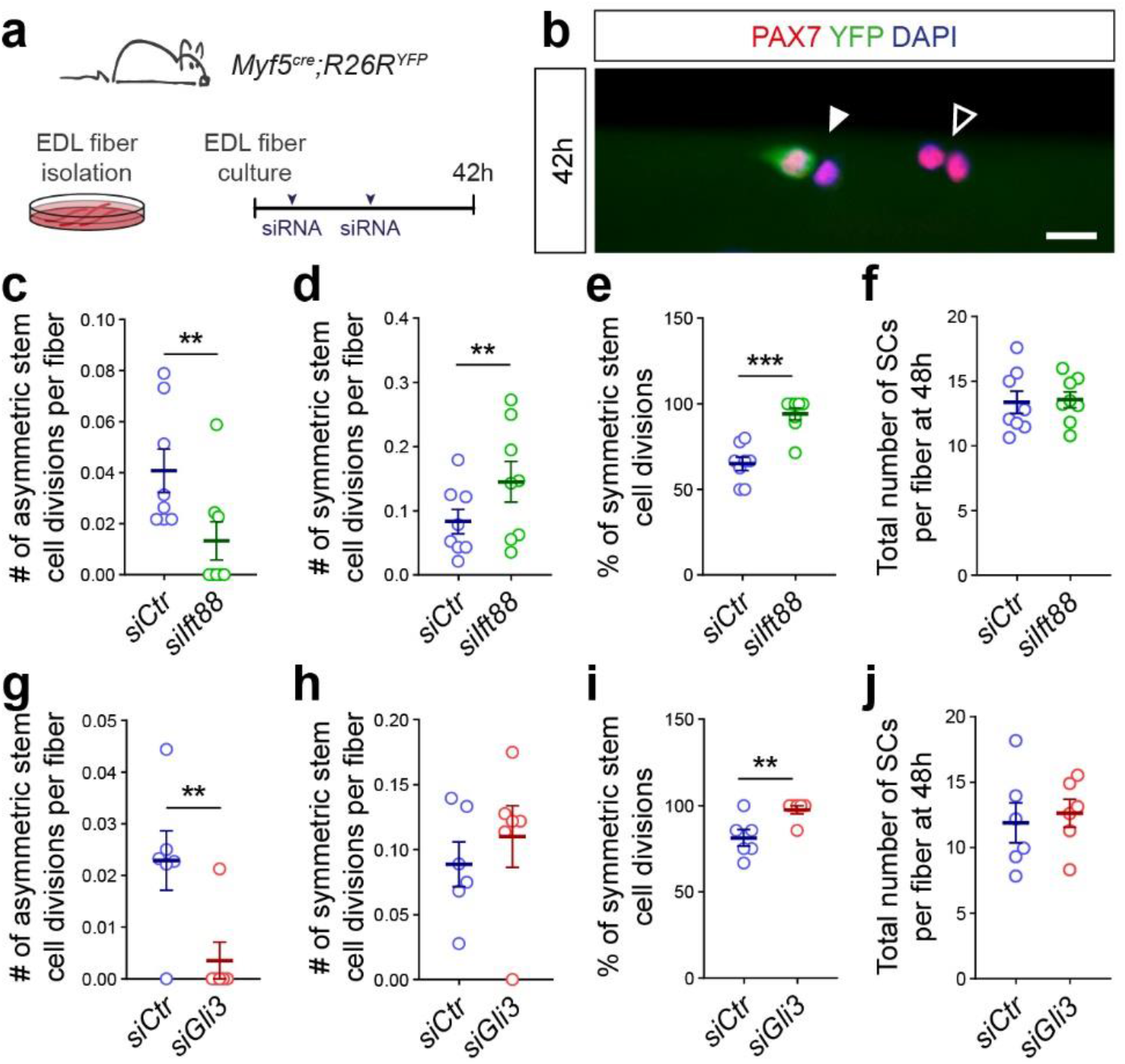
Primary cilia-mediated GLI3 processing regulates asymmetric division. **a**) Experimental design followed to knockdown *Gli3* or *Ift88* in satellite cells on cultured *Myf5^cre^:R26R^YFP^* single EDL myofibers. **b**) Representative immunofluorescence staining of PAX7 (red), YFP (green) and nuclei (DAPI, blue) showing an asymmetric (white arrow) and a symmetric (empty arrow) stem cell division on a *Myf5^cre^:R26R^YFP^* EDL myofiber cultured for 42h. **c**) Quantification of asymmetric stem cell doublets per fiber following treatment with siRNA against *Ift88 (silft88)* or with a non-target control *(siCtr)* (n = 8 males). **d**) Quantification of symmetric stem cell divisions per fiber following treatment with *silft88* or *siCtr* (n = 8 males). **e**) Proportion of symmetric stem cell divisions following treatment with *silft88* or *siCtr* (n = 8 males). **f**) Total number of PAX7^+^ satellite cells per fiber following treatment with siRNA against *Ift88 (silft88)* or with a non-target control *(siCtr)* (n = 8 males). **g**) Quantification of asymmetric stem cell divisions per fiber following treatment with siRNA against *Gli3 (siGli3)* or with *siCtr* (n = 6 males). **h**) Quantification of symmetric stem cell doublets per fiber following treatment with *siGli3* or *siCtr* (n = 6 males). **i**) Proportion of symmetric stem cell divisions following treatment with *siGli3* or *siCtr* (n = 6 males). **j**) Total number of PAX7^+^ satellite cells per fiber following treatment with siRNA against *Gli3 (siGli3)* or with a non-target control *(siCtr)*(n = 6 males). Scale bar, 10μm; Error bars, SEM; \*\**p* < 0.01; \*\*\**p* < 0.001.

To test the ability of satellite cells to proliferate following their first division, single EDL myofibers from *Gli3*^+/+^ and *Gli3*^Δ/Δ^ mice were cultured for 47h and incubated with 5-ethynyl-2’- deoxyuridine (EdU) 1h before fixation (**Supplementary Fig. 6a**). At 48h, *Gli3*^Δ/Δ^ satellite cells exhibit increased EdU incorporation along with an increase in the number of satellite cells per myofiber (**Fig. 7a, b; Supplementary Fig. 6b**). Analyzing *in vivo* EdU incorporation at 3 days postinjury further confirmed the enhanced ability of *Gli3*^Δ/Δ^ ASCs to proliferate (**Fig. 7c, d; Supplementary Fig. 6b)**. Single EDL myofibers from *Gli3*^+/+^ and *Gli3*^Δ/Δ^ mice were also cultured for 72h to assess their differentiation potential. At this time point, the proportion of cells expressing the myogenic differentiation factor MYOGENIN (MYOG^+^) was significantly decreased upon *Gli3* deletion (**Fig. 7e, f**), consistent with a reduction in generation of differentiation-competent progenitors due to decreased levels of asymmetric division^41^.

**Figure 7.**
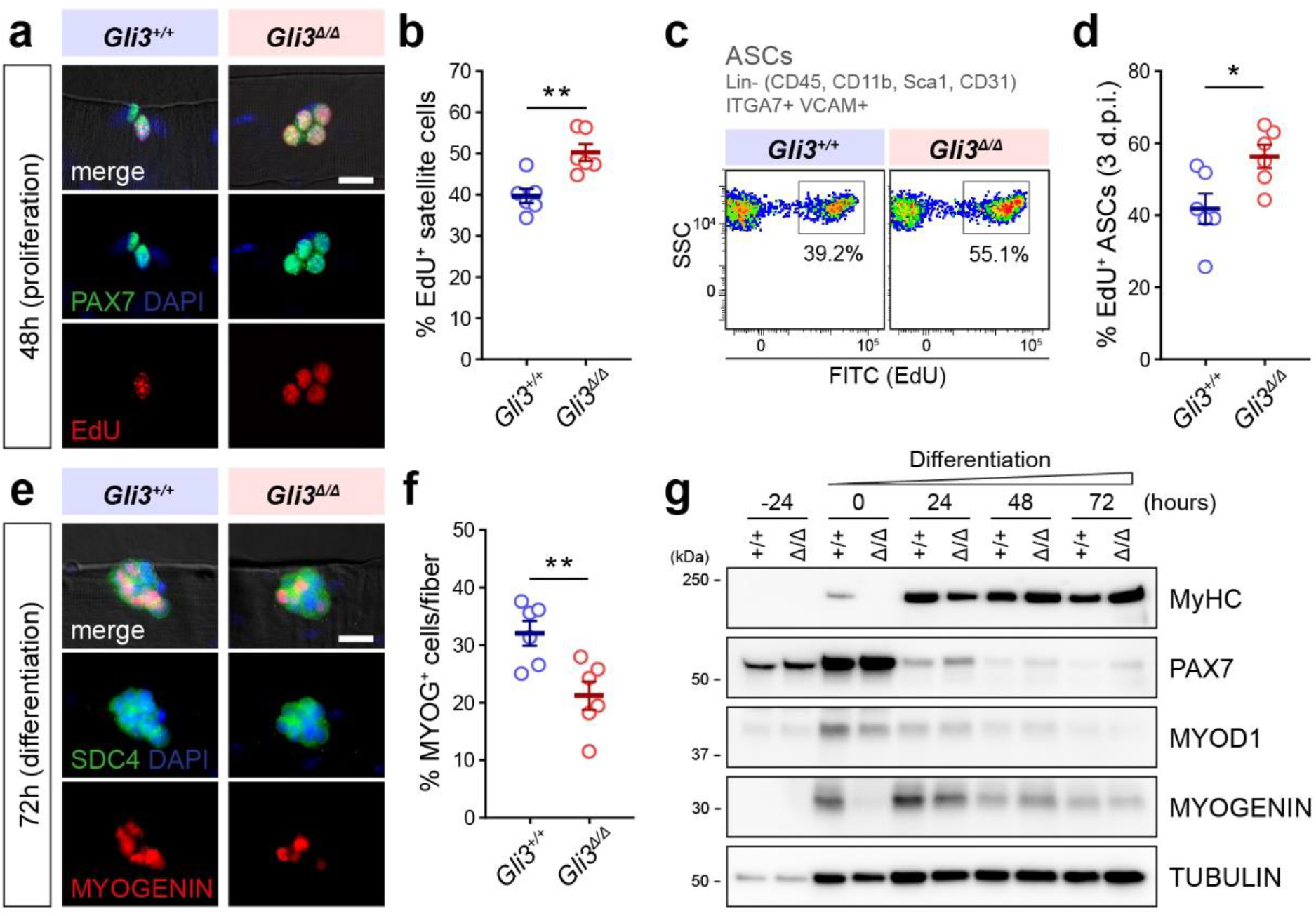
Gli3 deletion allows for the expansion of the satellite cell pool. **a**) Satellite cell immunostaining with PAX7 (green) and DAPI (nuclei, blue) showing EdU (red) incorporation by the PAX7^+^ cells after 1h EdU incubation. **b**) Proportion of *Gli3*^+/+^ and *Gli3*^Δ/Δ^ satellite cells that have incorporated EdU (EdU^+^). **c**) Representative flow cytometry plot of *Gli3*^+/+^ and *Gli3*^Δ/Δ^ activated satellite cells (ASCs) that have incorporated EdU. **d**) Proportion of EdU^+^*Gli3*^+/+^ and *Gli3*^Δ/Δ^ ASCs isolated from injured muscles at 3 days post-injury (d.p.i.). **e**) Representative immunofluorescence using anti-SYNDECAN-4 (SDC4, green) to label all satellite cells and MYOGENIN (MYOG, red) to mark the differentiated ones. DAPI stains the nuclei in blue. **f**) Quantification of the number of MYOG^+^ satellite cells per *Gli3*^+/+^ and *Gli3*^Δ/Δ^ myofiber after 72h of culture. **g**) Immunoblot analysis of MyHC, PAX7, MYOD1 and MYOGENIN from *Gli3*^+/+^ and *Gli3*^Δ/Δ^ myoblasts differentiated for 72h. TUBULIN is used as a loading control. Unless indicated otherwise, n = 6 (3 males and 3 females for each genotype); Scale bars, 10μm; Error bars, SEM; \**p* < 0.05; \*\**p* < 0.01

To determine whether *Gli3* deletion permanently impairs or transiently delays myogenic differentiation, we derived primary myoblasts from *Gli3*^+/+^ and *Gli3*^Δ/Δ^ satellite cells *in vitro*(**Supplementary Fig. 6a**). Conversely to QSCs and ASCs, RT-qPCR analysis revealed that *Gli3*^Δ/Δ^ primary myoblasts display increased expression of *Gli1* and *Ptch1* (**Supplementary Fig. 6c**), suggesting that GLI3R regulates the expression of its canonical target genes in myogenic progenitors. This is also consistent with our results showing that in wild-type primary myoblasts, the modulation of GLI3R level directly affects Gli1 and Ptch1 expression. (**Supplementary Fig. 2**). However, in *Gli3*^Δ/Δ^ myoblasts, SAG and FSK treatments failed to change the two GLI-target gene expression, excluding a role of GLI1 and GLI2 to compensate for the absence of GLI3 (**Supplementary Fig. 6d, e**). *Gli3*^Δ/Δ^ myoblasts exhibit decreased expression of the myogenic regulatory factors, MYOD1, MYOGENIN, and the Myosin Heavy Chain (MyHC), at the early steps of differentiation (0-24h), yet they express similar levels of MyHC at the later steps (48-72h) and even form bigger myotubes than *Gli3*^+/+^ cells (**Fig. 7g; Supplementary Fig. 6c, f**). This suggests that *Gli3*^Δ/Δ^ myoblast differentiation is delayed rather than impaired. Together, these results show that *Gli3* deletion increases satellite cell proliferation at the expense of early differentiation and delays, but does not prevent, terminal differentiation and fusion of myogenic progenitors.

### GLI3R controls the regenerative potential of muscle stem cells

To assess their regenerative potential, *Gli3*^+/+^ and *Gli3*^Δ/Δ^ mice were subjected to a CTX-induced injury in the *tibialis anterior* (TA) muscle (**Fig. 8a**). At 7 days post-injury (d.p.i.), we counted the numbers of self-renewing and differentiating cells expressing either PAX7 or MYOG, respectively. An increased number of PAX7^+^ SCs was observed in *Gli3*^Δ/Δ^ mice (**Fig. 8b, c; Supplementary Fig. 7a**), in accordance with increased satellite cell self-renewal (**Fig. 6**) and proliferation (**Fig. 7a-d**). The number of MYOG^+^ cells was slightly decreased compared to *Gli3*^+/+^ mice (**Supplementary Fig. 7b-c’**), correlating with our *ex vivo* and *in vitro* findings suggesting a delay in differentiation (**Fig. 7e-g**). Of note, the morphology of the nascent myofibers expressing DYSTROPHIN (DYS) is improved in *Gli3*^Δ/Δ^ mice, where interstitial space is reduced and myofibers are larger, suggesting that the delay in myogenic commitment does not impact the overall efficiency of regeneration (**Fig. 8b; Supplementary Fig. 7b**).

**Figure 8.**
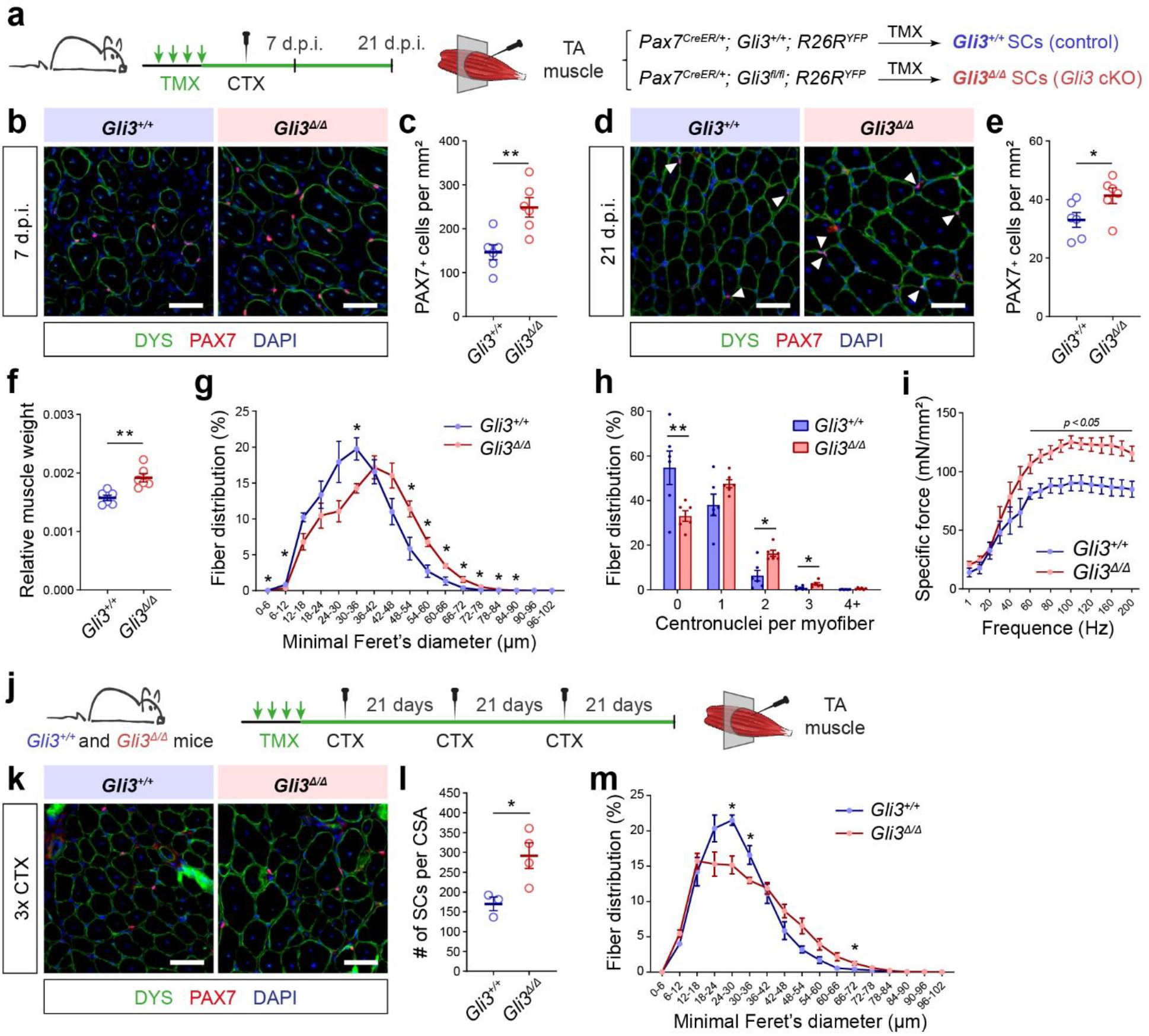
Gli3 deletion in satellite cells enhances muscle regeneration. **a**) Experimental design to analyze muscle regeneration of *Gli3*^+/+^ and *Gli3*^Δ/Δ^ mice. Muscle injury was induced by cardiotoxin (CTX) injections. **b**) Representative images of TA muscle sections at 7 days postinjury (d.p.i.). Satellite cells (SCs) are labelled with PAX7 (red). DYSTROPHIN (DYS, green) delineates the regenerating myofibers. **c**) Quantification of PAX7^+^ SCs per mm^2^ TA section of *Gli3*^+/+^ and *Gli3*^Δ/Δ^ mice at 7 d.p.i. **d**) Immunostaining of PAX7 and DYSTROPHIN on TA muscle sections at 21 d.p.i. **e**) Quantification of PAX7^+^ SCs per mm^2^ TA section, **f**) TA muscle weight normalized to total body weight, **g**) distribution of TA myofibers according to minimal Feret’s diameter and **h**) distribution of regenerated myofibers according to their number of centrally located nuclei (centronuclei) of *Gli3*^+/+^ and *Gli3*^Δ/Δ^ mice at 21 d.p.i. n = 6 (3 males and 3 females) **i**) Specific force of regenerated TA muscles from *Gli3*^+/+^ and *Gli3*^Δ/Δ^ mice at 21 d.p.i. (n = 5 males). **j**) Experimental design to analyze regeneration of *Gli3*^+/+^ and *Gli3*^Δ/Δ^ mice following three consecutive CTX-injuries in the TA muscle (3xCTX). **k**) Immunostaining of PAX7 and DYSTROPHIN on TA muscle sections at following 3xCTX. **l**) Quantification of PAX7^+^ satellite cells per cross-sectional area (CSA) of injured TA and **m**) distribution of regenerated TA myofiber according to minimal Feret’s diameter of 3xCTX *Gli3*^+/+^ and *Gli3*^Δ/Δ^ mice. DAPI stains nuclei (blue); Scale bars, 50μm; Error bars, SEM; \**p* < 0.05; \*\**p* < 0.01

The number of PAX7^+^ SCs was increased in *Gli3*^Δ/Δ^ mice at 21 d.p.i., further suggesting enhanced self-renewal and expansion of the SC pool (**Fig. 8d, e**). Additionally, there was a significant increase in muscle mass mainly due to myofiber hypertrophy (**Fig. 8f, g; Supplementary Fig. 7f-i’**) and increased numbers of centrally-located nuclei (**Fig. 8h**). This increase in myonuclear accretion suggests that, although there is a delay in terminal differentiation, the initial expansion of the satellite cell pool ultimately leads to a higher number of myogenic progenitors that eventually fuse into nascent and regenerating myofibers. Finally, *in situ* measurements of muscle force revealed that *Gli3*^Δ/Δ^ regenerated TA muscles were ~50% stronger than *Gli3*^+/+^ muscles (**Fig. 8i; Supplementary Fig. 7j**). Together our data show that *Gli3* deletion in satellite cells accelerates and improves muscle regeneration after CTX-induced trauma.

We also assessed the long-term regenerative capacity of the *Gli3*^Δ/Δ^ satellite cells by challenging the mice with repetitive muscle injuries (**Fig. 8j**). Strikingly, *Gli3*^Δ/Δ^ regenerated muscles are bigger, though some regions exhibit clear histological defects with fibrotic areas and tiny myofibers (**Supplementary Fig. 7k**). Even after three rounds of injury, *Gli3*^Δ/Δ^ mice maintain a higher number of self-renewing SCs (**Fig. 8k, l**), although their distribution throughout muscle section remains similar (**Supplementary Fig. 7m**). As observed following a single injury, *Gli3*^Δ/Δ^ mice exhibited increased muscle weight resulting from both myofiber hyperplasia and hypertrophy (**Fig. 8m; Supplementary Fig. 7l, n-p**), and regenerating myofibers contained a higher number of centrally located nuclei (**Supplementary Fig. 7q**).

This again suggests that during each regeneration cycle, the stem cell pool expansion provides increasing number of myogenic progenitors that ultimately fuse to restore the damaged myofiber. Altogether, our data demonstrate that GLI3R regulates the expansion of the SC pool during muscle regeneration, and controls their regenerative potential.

## DISCUSSION

Quiescent satellite cells lying in a non-cycling, dormant state have a primary cilium. Our study, as well as others^12,28,42^, demonstrates that muscle cells are transiently and dynamically ciliated as they progress through the myogenic lineage, and that the primary cilium reassembles in self-renewing stem cells. Here, we specifically identify the primary cilia-mediated processing of GLI3 as a downstream effector of the quiescent state, revealing a novel mechanism of muscle stem cell regulation. Indeed, we find that satellite cell-specific depletion of GLI3 induces G_Alert_ in QSCs. Moreover, we show that *Gli3* deletion promotes stem cell expansion and enhances regenerative and engraftment potential, providing a proof-of-principle for prospective therapeutic applications of our findings.

Our results show that GLI3R colocalizes with PKA at the basal bodies of QSCs. Many studies have delineated the signaling networks that maintain satellite cell quiescence^2,4,5,43-48^. One such pathway, the NOTCH-COLV-Calcitonin receptor (CALCR) cascade, acts through the PKA pathway, which also promotes GLI3 phosphorylation and processing into the repressor form^4,46,48^. Therefore, one can hypothesize that CALCR-dependent maintenance of G_0_ in QSCs is mediated through GLI3 phosphorylation. Upon muscle damage, disruption of the stem cell niche downregulates NOTCH signaling and cAMP-PKA activity^4,46^, potentially abrogating GLI3 phosphorylation and processing, which could then promote satellite cell activation independent of Hedgehog ligand-receptor binding. It is highly likely that interaction between signaling networks, such as this, underlies the fine-tuned control of satellite cell activation during regeneration, and further investigations into such interplay will be essential for the development of our understanding of satellite cell biology.

While our study shows that GLI3 processing is required to maintain a dormant quiescent state, *Gli3* deletion does not lead to autonomous activation in satellite cells but instead induces an “alert” quiescence. The G_Alert_ state was first described in quiescent stem cells subjected to systemic exposure of HGFA released from a distant muscle injury^6,9^. Interestingly, fully reduced HMGB1 released from a bone injury induces G_Alert_ in SCs through CXCR4 signaling^10^, suggesting that several signaling pathways can induce the G_Alert_ state in stem cells. In addition, Der Vartanian et al. demonstrated that a subpopulation of satellite cells is protected from dioxin pollutant through a cell intrinsic, mTORC1-dependent G_Alert_ response^11^, demonstrating that factors released upon injury are not mandatory to induce G_Alert_. Here, we find that the intrinsic loss of GLI3R is sufficient to induce G_Alert_ in satellite cells in the absence of any systemic or extrinsic cues. In wild-type condition, processing of GLI3 is likely to be responsive to both extrinsic and intrinsic cues, such as the CALCR discussed above. Hence, we propose that cilia-mediated GLI3 processing controls muscle stem cell quiescence and regulates the first step of entering G_Alert_ in a physiological context.

*Gli3*^Δ/Δ^ resting muscle exhibit a slight increase in the number of satellite cells two weeks post-tamoxifen treatment. However, this number remains stable over time, and our results indicate that *Gli3*^Δ/Δ^ satellite cells return to and maintain quiescence under homeostatic conditions. Satellite cells contribute to uninjured myofiber homeostasis in adulthood^49,50^. Since *Gli3* deletion leads to enhanced cell proliferation and self-renewal, we speculate that the initial depletion of *Gli3* triggers a transient entry into the cell cycle with a preference for self-renewal, increasing the total number of satellite cells before homeostatic quiescence is re-asserted.

Intriguingly, our RNA-sequencing data indicates that deleting *Gli3* from the QSCs and ASCs does not significantly impact the expression of the putative GLI-target genes of the canonical Hedgehog pathway, notably *Gli1* or *Ptch1.* Instead, loss of GLI3R induces the activation of mTORC1 signaling. Interestingly, the upregulation of the GLI transcription factors in ischemia/reperfusion injury protects muscle tissue through activation of AKT/mTOR/p70S6K signaling^51^. Several studies have highlighted the existence of a GLI-mediated mTORC1 activation^52,53^, where GLI transcription factors regulate positively mTORC1 signaling by downregulation of negative or upregulation of positive mTORC1 mediators^54^. Both *Gli3*^+/+^ and *Gli3*^Δ/Δ^ QSCs and ASCs display decreased expression of *Deptor*, a negative regulator of mTORC1. Although *Deptor* is not a direct target of GLI3, downregulation of *Deptor* in *Gli3*^Δ/Δ^ satellite cells is consistent with increased mTORC1 activity and S6 phosphorylation, consequently leading to G_Alert_ transition in *Gli3*^Δ/Δ^ QSCs as well as enhanced proliferation in *Gli3*^Δ/Δ^ ASCs^6,38^.

Activated satellite cells undergo either asymmetric or symmetric division, giving rise to one committed muscle progenitor and one self-renewing stem cells, or to two self-renewing stem cells, respectively^7,55^. Stimulating symmetric stem cell expansion is associated with improved muscle regeneration^39,40,56^. Interestingly, GSEA revealed that *Gli3*^Δ/Δ^ satellite cells display inhibition of the JAK/STAT signaling, which knockdown or pharmacological inhibition has been shown to favor satellite cell expansion, homing and regenerative capacity, overall improving skeletal muscle repair^40,56^. Thus, the decreased activity of JAK/STAT signaling in *Gli3*^Δ/Δ^ satellite cells is consistent with their enhanced ability to self-renew and regenerate. As well, GO analysis on the 629 downregulated genes highlighted biological process terms mainly related to cell adhesion, supporting the idea that *Gli3*^Δ/Δ^ ASCs are less adhesive to their environment during regeneration, allowing more efficient migration and muscle tissue repair^2,57^. Corroborating this result, GSEA revealed the inhibition of cell-cell interaction mechanisms (Apical junction/surface, Epithelial-mesenchymal transition). Thus, increased self-renewal, decreased adhesion and enhanced proliferation overall improves the regenerative capacity of *Gli3*^Δ/Δ^‘alert’ satellite cells.

Nevertheless, sustained cell proliferation can compromise myogenic differentiation and impair muscle regeneration^56^. As previously shown^58^, the absence of GLI3 in myogenic progenitors increases their proliferation at the expense of early differentiation. However, while *Gli3* deletion in differentiated muscle cells delays the overall muscle regeneration^58^, *Gli3* deletion in satellite cells leads to opposite. *Gli3*^Δ/Δ^ satellite cells form larger myotubes upon differentiation *in vitro* and promote myofiber hypertrophy *in vivo*. Likely, this discrepancy is the result of the expansion of satellite cells at the early stages of regeneration/myogenesis, providing a larger pool of progenitor myoblasts that eventually, after a delay, fuse into myofibers or myotubes. The expansion is mediated both by the G_Alert_ state, conferring accelerated proliferative properties to satellite cells, and the increased propensity for self-renewal divisions. Ultimately, the initial expansion compensates for the delay in myogenic commitment, leading to better repair in *Gli3*^Δ/Δ^ mice. As we and others showed^42^, GLI3 is down-regulated during the later stages of myogenic differentiation, excluding a potential role for GLI3 after satellite cell commitment. This suggests that other signals emanating from the regenerating muscle are sufficient to induce myogenesis^59^ and this response does not appear to be impaired in *Gli3*^Δ/Δ^ satellite cells.

Different studies have explored the beneficial effects of activating canonical Hedgehog signaling to induce myogenic progenitor proliferation and differentiation during muscle repair^51,60-65^. Our results show that satellite cells do not activate canonical Hedgehog signaling within the first 3 days following an acute injury. While deleting *Gli3* from satellite cells does not activate the expression of the canonical GLI-target genes, increased expression of *Gli1* and *Ptch1* was observed in SAG-treated myoblasts and *Gli3*^Δ/Δ^ myogenic progenitors, suggesting that they can respond to Hedgehog ligands. At day 5 following CTX-induced acute injury, when the population of myogenic progenitors is predominant, DHH is robustly expressed by the Schwann cells^27^. Thus, the induction of DHH might contribute to the proliferation and differentiation of the myogenic progenitors. Our results imply that GLI3R has pleiotropic roles during muscle stem cell progression through the myogenic lineage: controlling satellite cell quiescence and activation independently of its canonical target genes while repressing Hedgehog signaling target genes in myogenic progenitors to regulate their proliferation and differentiation.

Our results demonstrate that the primary cilium-mediated control of GLI3 processing regulates muscle stem cell fate and that loss of GLI3R from activated satellite cells and myoblasts results in increased proliferation and self-renewal. However, studies regarding the ablation of primary cilia from muscle cells have led to different and somewhat conflicting results^12,42^. In the C2C12 myogenic cell line, *Ift88* knockdown results in increased proliferation at the expense of differentiation^42^. Conversely, drug-mediated cilia disassembly does not affect satellite cell proliferation and differentiation but impairs self-renewal^12^. This discrepancy is likely related to the experimental settings and differences in the behavior of myogenic progenitors and satellite cells, as we showed. As well, these data suggest that the primary cilium has broader signaling function in muscle cells than the only regulation of GLI3 proteolytic processing.

Although we largely observed beneficial effects of GLI3 depletion, we cannot exclude the possibility that long-term loss of GLI3 repressor function can have negative consequences, as observed in other knockout models^66^. *Gli3*^Δ/Δ^ regenerated muscles following a triple injury exhibit signs of histological defects, suggesting that permanent GLI3R depletion may eventually lead to regenerative deficits over time. Along these lines, given the importance of quiescence regulation in preventing precocious activation and maintaining the satellite cell pool over an organism’s lifetime, one could speculate that the negative consequences of *Gli3*^Δ/Δ^ would not be observed until much later ages. As well, the persistence of a G_Alert_ state in uninjured homeostasis represents an inefficient use of energetic and substrate resources within muscle, which could be detrimental in contexts of metabolic scarcity. Therefore, translational studies should aim to determine the benefits of transient GLI3 depletion or the pharmacological inhibition of its proteolytic cleavage for the development of stem cell-based therapeutic strategies in regenerative medicine^44,66-68^.

Our findings represent a seminal advancement in our understanding of the molecular regulation of adult muscle stem cell function. This study establishes, for the first time, that primary cilia-mediated GLI3 processing controls the transition from G_0_-to-G_Alert_, as well as the fate-determinate division that follows satellite cell activation. As a result, GLI3 processing within the primary cilia directly impacts the regenerative capacity of stem cells. Further studies are needed to understand how endogenous signaling events or mechanosensing by the primary cilia can control GLI3 processing in response to injury. Also, the downstream signaling events that mediate GLI3R activity remain to be determined. Finally, pharmacological methods of manipulating GLI3 processing warrant investigation for their potential use in muscle stem cell-based therapies.

## METHODS

### Mouse strains and animal care

The following mouse lines were used in this study: *Gli3^fl/fl^* mice69, Pax7CreERT2/+ mice70 referred as *Pax7^CE/+^* in the text, *Myf5^Cre^* mice^71^, *R26R^EYFP^* mice^72^ referred as *R26R^YFP^, Pax7nGFP* mice^73^, C57BL/10ScSn-*Dmd^mdx^*/J mice (homozygous *Dmd^mdx^* females and hemizygous *Dmd^mdx^* males) referred as MDX in the manuscript. All the mice used in this study were males and females, from 2 to 10 month-old, with mixed genetic background (129SV and C57BL/6). Mice were sex and age-matched in all experiments. Housing, husbandry and all experimental protocols for mice used in this study were performed in accordance with the guidelines established by the University of Ottawa Animal Care Committee, which is based on the guidelines of the Canadian Council on Animal Care (CCAC). Protocols were approved by Animal Research Ethics Board (AREB) at the University of Ottawa.

### Tamoxifen treatment and muscle injury

Both *Pax7^CE/+^:Gli3^+/+^:R26R^YFP^* and *Pax7^CE/+^:Gli3^mF^R26R^YFP^* mice were injected intraperitoneally with 100μl of a 20mg.ml^−1^ tamoxifen solution (TMX, Sigma T5648) dissolved in corn oil for 4 consecutive days, and then they were maintained on a diet containing tamoxifen (500mg TMX per kg diet, Teklad, Envigo). Muscle injury was induced by intramuscular injections of cardiotoxin (CTX; 10μM; Latoxan). Mice were administered buprenorphine and then, anesthetized by isofluorane inhalation. For histological analysis, 50μl of was injected into *tibialis anterior* (TA) muscle. For flow cytometry and FACS analysis, both TA and *gastrocnemius* muscles were injected with 40μl and 80μl of CTX, respectively.

### Muscle fixation and histological analysis

Mice were euthanized and TA muscles were harvested, weighed and embedded in OCT and frozen in liquid nitrogen-cooled isopentane. For GFP visualization, mice were euthanized with a sodium pentobarbital solution (Euthanyl) and perfused with PBS before fixation with 4% paraformaldehyde (PFA) solution in PBS. Muscles were excised, further fixed into 1% PFA/PBS for 16h, then transferred into ascending sucrose gradients (15%, 30% in PBS), and finally embedded and frozen in OCT. Embedded muscles were transversely sectioned at 10μm thickness. Sections were post-fixed in 4% PFA/PBS 10min at room temperature, permeabilized in 0.1M glycine, 0.1% Triton X-100 in PBS, and blocked in 5% goat serum, 2% BSA in PBS supplemented with M.O.M. Blocking reagent (Vector Laboratories). Then, sections were incubated with primary antibodies as described in “Immunostaining on cells, myofibers and sections”. Muscle cross-sections stained with anti-Laminin and anti-Dystrophin and counterstained with DAPI were analyzed for fiber counting and minimum Feret’s diameter using SMASH^74^, and centronuclei per myofiber were quantified using MuscleJ^75^.

### Flow cytometry and Fluorescence-activated cell sorting (FACS)

Quiescent satellite cells were obtained from uninjured hindlimb muscles, while activated satellite cells were obtained from CTX-injured *tibialis anterior* and *gastrocnemius* muscles 3 days after the induced-injury. Dissected muscles were minced in collagenase/dispase solution followed by dissociation using the gentleMACS Octo Dissociator with Heaters (Miltenyi Biotec). Satellite cells were sorted by gating a mononuclear cell population of α7-INTEGRIN^+^, CD34^+^, CD31^neg^, CD45^neg^,

SCA1^neg^, CD11b^neg^ (quiescent satellite cells) or α7-INTEGRIN^+^, VCAM1^+^, CD31^neg^, CD45^neg^, SCA1^neg^, CD11b^neg^ (activated satellite cells) using a MoFlo XDP cell sorter (Beckman Coulter). Gating strategy is shown in **Supplementary Fig. 8**. Flow cytometry analyses (PyroninY, MitoTracker) were performed on a BD LSRFortessa cell analyzer (BD Biosciences). The list of antibodies is available in **Supplementary Table 1**.

### Cell size measurement

Quiescent and activated satellite cells were sorted based for α7-INTEGRIN^+^, VCAM1^+^, CD31^neg^, CD45^neg^, SCA1^neg^, CD11b^neg^. Single cell brightfield images were captured on the Amnis ImageStream XMk II and analyzed on the IDEAS Software.

### PyroninY and MitoTracker staining

40nM PyroninY (Santa Cruz) and 40nM MitoTracker Deep Red (ThermoFisher) were added to the muscle digests and incubated 30min at 37°C in a water bath. Then, muscle digests were washed and stained with the antibodies for gating the satellite cell population (α7-INTEGRIN^+^, VCAM1^+^, CD31^neg^, CD45^neg^, SCA1^neg^, CD11b^neg^). Gating strategy is shown in **Supplementary Fig. 5b**.

### In vivo EdU incorporation assay

Both tamoxifen-treated *Pax7^CE/+^:Gli3^+/+^:R26R^YFP^* and *Pax7^CE/+^;Gli3^fl/fl^;R26R^YFP^* mice were subjected to CTX-induced muscle injury in both TA and GA muscles. 3 days post-injury, mice were injected intraperitoneally with 10μl per gram of body weight of a 10mM EdU solution 3 hours before sacrifice. Then, muscles were collected and digested as described in ‘Flow cytometry and Fluorescence-activated cell sorting’ section.

### Satellite cell transplantation and engraftment assay

Host MDX mice were anesthetized with isofluorane and hindlimbs were irradiated with 16Gy X-rays delivered at 0.71Gy/min with an X-RAD 320 (Precision X-Ray) biological irradiator. The following day, irradiated MDX mice were implanted subcutaneously with osmotic pumps (Alzet) delivering FK-506 immunosuppressant (LC Laboratories) at 2.5mg/kg/day. Two days after, donor satellite cells from tamoxifen-treated *Pax7^CE/+^:Gli3^+/+^:R26R^YFP^* and *Pax7^CE+^;Gli3^fl/fl^;R26R^YFP^* mice were FACS-isolated based on FSC/SSC, lineage negative selection (CD31, CD11b, CD45, SCA1), and positive selection (α7-INTEGRIN, CD34). Donor satellite cells were washed with PBS and resuspended in 0.9% NaCl solution at a concentration of 10^5^ cells per 10μl prior to engraftment into the TA muscle of irradiated and immunosuppressed MDX mice. Transplanted TA muscles were collected two weeks after transplantation for measurement of satellite cell engraftment.

### EDL myofiber isolation, siRNA transfection and EdU treatment

Myofibers were isolated from *extensor digitorum longus* (EDL) muscles following the previously described protocol^76^. Briefly, EDL muscles were dissected from tendon-to-tendon and incubated for 1h in DMEM (Gibco) containing 0.25% collagenase I (Worthington). Single EDL myofibers were isolated by gentle muscle trituration and washed in DMEM. EDL myofibers were finally cultured in DMEM supplemented with 20% fetal bovine serum, 1% chick embryo extract and 1% penicillin/streptomycin. Myofibers were fixed at the desired time points using either PFA4%/PBS or ice-cold methanol for immunostaining analysis.

Single EDL myofibers were transfected with *Gli3* siRNA (TriFECTa DsiRNA Kit mouse Gli3, mm.Ri.Gli3.13), *Ift88* siRNA (TriFECTa DsiRNA Kit mouse Ift88, mm.Ri.Ift88.13) or scramble negative control siRNA, at a final concentration of 5nM using Lipofectamine RNAiMAX (Invitrogen) according to the manufacturer’s instructions. siRNA transfection was performed twice at 4h and 16h post-culture and 6h after the second transfection, growth medium was renewed.

For EdU treatment, freshly isolated EDL myofibers were treated for 24h with 20μM EdU (ThermoFisher) to analyze satellite cell activation. For cell proliferation analysis, 48h-cultured myofibers were treated with 20μM EdU for 1h before fixation.

### Myoblast isolation, culture and treatments

8-16 week-old mice were used to derive primary myoblasts by magnetic cell separation (MACS)^77^. Muscle dissociation and cell filtration was performed following the same protocol described for Flow cytometry and FACS. First, negative lineage selection was performed with biotin-conjugated lineage antibodies (CD11b, SCA1, CD45, CD31), followed by incubation with streptavidin microbeads. Then, satellite cell-derived myoblasts were purified using biotin-conjugated anti-α7- INTEGRIN antibody. Myoblasts were cultured on collagen-coated dishes in Ham’s F10 medium (Wisent) supplemented with 20% FBS, 1% penicillin/streptomycin, and 5ng.ml^−1^ of basic FGF (Millipore). Differentiation was induced in Ham’s F10:DMEM 1:1 supplemented with 5% horse serum, and 1% penicillin/streptomycin.

For GLI3 proteolytic processing analysis, primary myoblasts were treated with either 200mM SAG (Smoothened agonist, R&D Systems) or 25μM FSK (Forskolin, R&D Systems) for 24 hours. Equivalent amounts of DMSO were added to the control conditions.

### Reserve cell isolation

Primary myoblasts were derived from 8-16 week-old *Pax7-nGFP* mice and cultured as described above. Myoblasts were differentiated for 3 days. Then, total cells were trypsinized, washed in PBS and resuspended in FACS buffer. ‘Reserve’ cells were sorted by gating a mononuclear GFP^+^ cell population using the MoFlo XDP cell sorter (Beckman Coulter).

### RNA extraction and quantitative PCR

Total RNA was extracted from primary myoblasts using the Nucleospin RNA II kit (Macherey-Nagel) and from satellite cells using the ARCTURUS Picopure RNA extraction kit (ThermoFisher), according to the manufacturers’ instructions. Reverse transcription was performed using SuperScript III Reverse Transcriptase (Invitrogen). Gene expression was assessed with iQ SYBR Green Supermix (Bio-Rad) and analysis was performed using the 2^−ΔΔCt^ method. RT-qPCR were normalized to the housekeeping genes *Ppia*. A list of primers is available in **Supplementary Table 2**.

### RNA-sequencing and gene expression analysis

12 samples were used for RNA-sequencing analysis: 3 samples of quiescent satellite cells and 3 samples of activated satellite cells for each genotype *(Gli3^+/+^* and *Gli3^Δ/Δ^).* For each sample of quiescent satellite cells, all hindlimb muscles from 2-3 mice were combined. Activated satellite cells were isolated from injured TA and *gastrocnemius* muscles at day 3 post-CTX injections, and each sample corresponded to one mouse.

Library construction was performed with 20ng of input total RNA using the NEBNext Ultra II Directional RNA Library Prep Kit for Illumina - polyA mRNA workflow (New England Biolabs). The libraries were sequenced with a NextSeq 500 High Output 75 cycle kit (Illumina). RNA-seq reads were mapped to transcripts from GRCm38_GENCODE.vM19 using salmon v0.13.1^78^. Data were loaded into R using the tximport library and the gene/count matrix was filtered to retain only genes with five or more mapped reads in two or more samples. Differential expression was assessed using DESeq2^79^. PCA was performed using the DESeq2 plotPCA function and rlog-transformed count data. Pearson correlation between means of CPM normalized expression for each replicate group was calculated. Expression differences were calculated using the lfcShrink function, applying the apeglm method (v1.6.0)^80^. Gene Ontology (GO) analysis and Gene set enrichment analysis (GSEA) were performed using goseq (v1.40.0) and fgsea (v1.14.0) R packages, respectively, on the significantly upregulated and downregulated genes (cut-off of 0.05 and absolute fold changes greater than or equal to 1.2) from *Gli3*^+/+^ and *Gli3*^Δ/Δ^ QSCs and ASCs.

### Western blotting

Whole cell proteins were extracted in lysis buffer (150mM NaCl, 25mM Tris pH7.5, 1% NP-40, 0.5% sodium deoxycholate, 0.1% SDS) supplemented with inhibitors of proteases (Roche) and phosphatases (Sigma). Equal amounts of proteins were resolved on SDS-PAGE 4-12% (Bio-Rad) and transferred onto PVDF membranes. Membranes were blocked using 5% non-fat dry milk in TBS-Tween 0.1% (TBST) for 1h at room and probed with primary antibodies overnight at 4°C. The list of antibodies is available in **Supplementary Table 1**. After 4 washes in TBST, membranes were incubated 1h with HRP-conjugated secondary antibodies at 1:5,000 (Bio-Rad). After 4 more washes, immunoblots were developed by enhanced chemiluminescence. When required, PVDF membranes were stripped in 62.5mM Tris HCl, pH6.8, 2% SDS and 0.8%β-mercaptoethanol.

### Immunostaining on cells, myofibers and sections

Following fixation in PFA 2%/PBS, cells and myofibers were washed 2 times in PBS, permeabilized in 0.1M Glycine, 0.1% Triton X-100 in PBS for 10min. For EdU staining, samples were stained using the Click-iT EdU Alexa Fluor 647 Imaging kit (ThermoFisher), according to the manufacturer’s instructions. Then, cells and myofibers were blocked in 5% horse serum, 2% BSA, 0.1 % Triton X-100 in PBS for at least 1h, and incubated with primary antibodies overnight at 4°C. The list of antibodies is available in **Supplementary Table 1**. Samples were washed 3 times in PBS, incubated 1h at room temperature with Alexa Fluor-conjugated secondary antibodies at 1:1,000 (ThermoFisher), washed 3 times in PBS and counterstained with DAPI at 1 μg.mL^−1^ in PBS before mounting.

For GLI3/acαTUB/PAX7 co-staining, cells and myofibers were first incubated with anti-GLI3 primary antibody and then, Alexa Fluor donkey anti-goat secondary antibody. Following 2 washes in PBS-Tween 0.1% and PBS, cells and myofibers were incubated with anti-acαTUB and anti-PAX7 primary antibodies followed by Alexa Fluor goat anti-mouse IgG2b and goat anti-mouse IgG1 secondary antibodies.

γTUB immunostaining required ice-cold methanol fixation. Briefly, cells and myofibers were rinsed two times in PBS, incubated 30min in ice-cold MeOH at −20°C and rehydrated in successive washes of 50% MeOH/50% PBS, 30% MeOH/70% PBS and PBS before being incubated in blocking buffer.

Full muscle section pictures were taken on a Zeiss Axio Observer.D1 inverted microscope equipped with an EC Plan-Neofluar 10×/0.3 Ph1 M27 objective and stitched together using Fiji software (http://fiji.sc/Fij). Other immunofluorescence pictures were taken with a Zeiss Axio Observer.D1 inverted microscope equipped with either a Plan-Apochromat 20×/0.8 M27 objective or a Plan-Apochromat 63×/1.4.Oil DIC M27 objective. For GLI3 localization, images were taken with a confocal Zeiss LSM 880 AiryScan inverted microscope quipped with a Plan-Apochromat 63×/1.4.Oil DIC M27 objective. Images were processed and analyzed with Zen and FIJI software.

### In situ force measurement

Muscle force measurements were performed on an Aurora Scientific 300C-LR-FP dual mode muscle lever system equipped with a 1N force transducer and 1cm lever arm. Electrical stimulation was performed using monopolar needle electrodes attached to an Aurora Scientific 701C High-Power, Bi-Phase Stimulator. Force transducers were calibrated prior to the study using precision weights. Mice were anesthetized using isoflurane inhalation (2% isoflurane, 1L/min) until recumbent and non-reflexive to pressure on the paw and positioned on a heated pad to maintain their body temperature at 37°C throughout the procedure. Mice were positioned supine and hindlimb were shaved. A small incision was made above the hallux and the foot was partially degloved to expose the distal insertion of the tibialis anterior tendon up to the tibialis anterior muscle. The cruciate crural ligament was severed to release the *tibialis anterior* (TA) tendon from the foot.

A pre-tied loop of waxed 3.5 metric suture was attached to the TA tendon using a series of double thumb knots above, below and through the loop. The suture was secured to the tendon using minimal amounts of cyanoacrylate glue. The skin of the hindlimb was removed up to mid *vastus lateralis* to expose the TA and the kneecap. The fascia of the TA was cut using spring scissors. The distal insertion of the TA tendon was severed and the TA was gently lifted to release it from the *extensor digitorum longus* (EDL) muscle and connective tissue. Muscles were kept from drying using physiological saline. The measured hindlimb was secured between the limb clamp and the stage using a 40mm long 27g needle inserted through the epiphysis of the femur immediately proximal the kneecap and directly into a receiving hole in the stage. Clamping was verified by observing no movement of the kneecap following manipulation of the foot and the needle was secured by a hand screw. The pre-tied loop was attached to the hook on the force transducer lever arm and maintained without tension.

Two monopolar needle electrodes were positioned adjacent to the tibial nerve proximal to the kneecap and distal the kneecap adjacent the EDL muscle. The transducer was retracted to maintain 20mN of measured tension for an initial 15 minute stretching period with 100ms trains of 0.3ms, 5V supramaximal voltage pulses at 1Hz stimulation every 100 seconds. Following stretching, muscles were maintained at 20mN tension and tetanic contractions were measured every 100 seconds following 200ms trains of 0.3ms, 5V supramaximal voltage pulses at serial frequencies from 1Hz to 200Hz. Maximal force was defined by the difference in maximal force measured during stimulation to that of the tension immediately prior stimulation.

### Statistical analysis

No statistical method was used to predetermine sample size. Statistical evaluation was performed using the Student’s t-test tests to calculate differences between two groups and either one-way or two-way ANOVA with post hoc test for multiple comparisons (Graphpad Prism®, **Data Source file**). The number of independent experimental replications is reported in each corresponding figure legend. Data are presented as mean ± S.E.M. and *p*-value < 0.05 was considered as statistically significant. Throughout the manuscript, level of significance is indicated as follows: \**p*≤ 0.05, \*\**p*≤ 0.01, \*\*\**p*≤ 0.001.

## ACKNOWLEDGMENTS

The authors thank Dr. Valerie Wallace for providing the *Gli3* floxed mice, Jennifer Ritchie for animal husbandry, Fernando Ortiz for FACS, Caroline Vergette from StemCore Laboratories, Gareth Palidwor from Bioinformatics Core, Hani Jrade and Damian Carragher for helping with the ImageStream flow cytometry, Alireza Ghasemizadeh for helping with confocal imaging, Hong Ming, Ricardo Carmona and Pascale Muller for technical assistance and Sandy Martino for administrative assistance. C.E.B. was supported by postdoctoral fellowships from the Ontario Institute for Regenerative Medicine (OIRM) and the French Muscular Dystrophy Association (AFM)-Téléthon and is now supported by a postdoctoral fellowship from the Fondation pour la Recherche Médicale [FRM, ARF201909009155]. A.Y.T.L. is supported by a postdoctoral fellowship from OIRM. P.F. was supported by a doctoral fellowship from the Canadian Institutes of Health Research (CIHR). Studies from the F.L.G. lab were supported by grants from the Agence Nationale pour la Recherche (ANR): Myofuse project [ANR-19-CE13-0016-03] and Myofibrosis project [ANR-19-CE14-0008-02], and from the European Joint Programme on Rare Diseases (EJP RD): MYOCITY project. M.A.R. holds a Canada Research Chair in Molecular Genetics. These studies were carried out with the support of grants from the US National Institutes for Health [R01AR044031], the Canadian Institutes of Health Research [FDN-148387], and the Stem Cell Network.

## AUTHOR CONTRIBUTIONS

C.E.B. and M.A.R. conceived the project. C.E.B., M.-C.S. and A.Y.T.L. designed methods. C.E.B., M.-C.S., A.Y.T.L. and D.H. conducted experiments and analyzed results. W.J. and F.L.G. performed *in silico* analysis of RNA-sequencing data. P.F. performed force measurement and analysis. M.R. performed satellite cell transplantation. C.E.B. wrote the original draft with input from co-authors. C.E.B. and M.A.R. edited the final manuscript. M.A.R. and F.L.G. provided financial support.

## SUPPLEMENTARY FIGURES

**Supplementary Figure 1.**
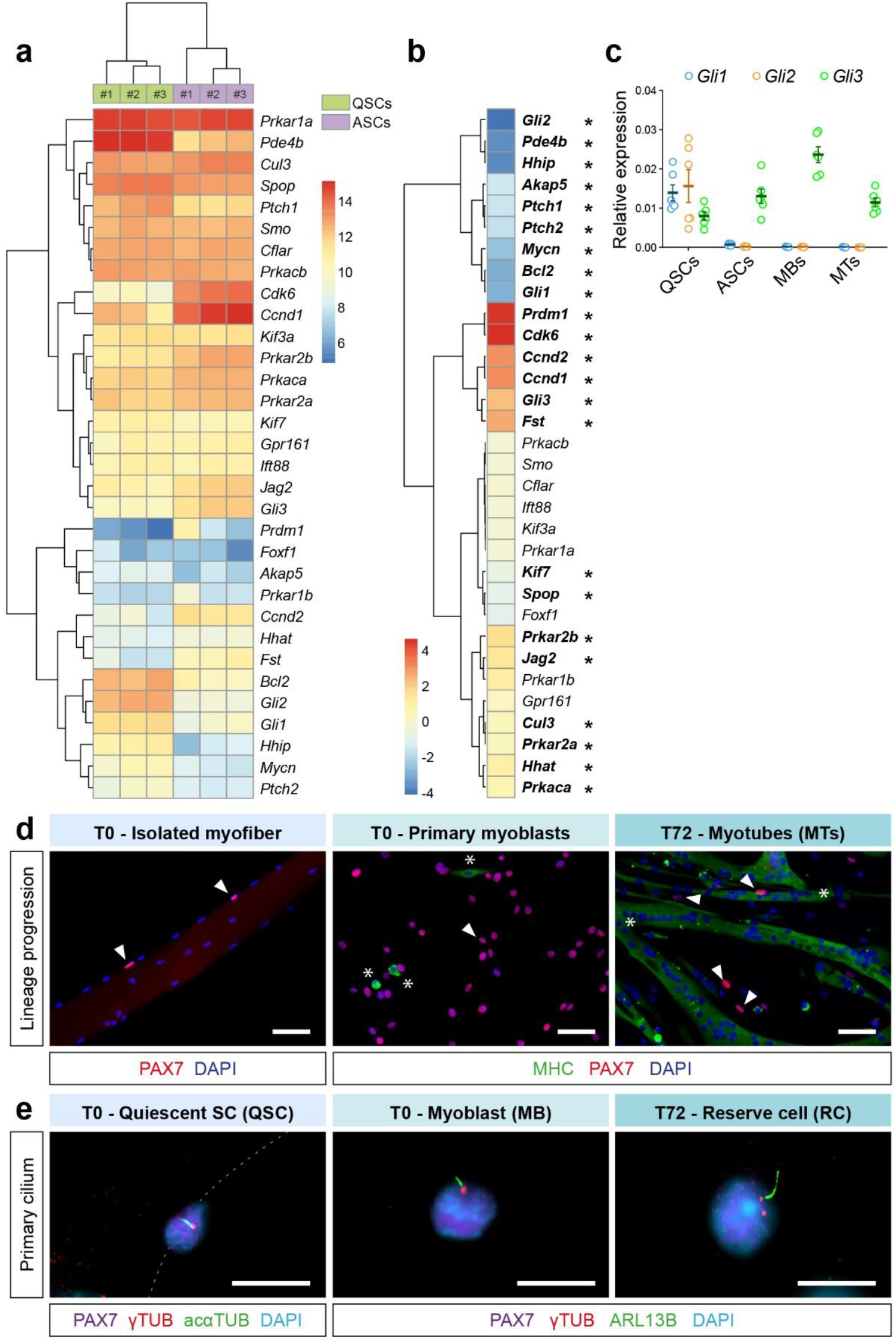
Gli3 is expressed in muscle cells throughout their lineage progression. **a**) Heatmap from normalized and log2 transformed expression matrix of components and target genes of canonical Hedgehog signaling in quiescent (QSCs) and activated satellite cells (ASCs) (n = 3 males). **b**) Heatmap showing fold change values of canonical Hedgehog signaling genes in ASCs compared to QSCs. Significantly up- and down-regulated genes are written in bold with a star that indicates *p* adjusted values (padj) < 0.05. **c**) Expression of *Gli1, Gli2* and *Gli3* determined by RT-qPCR and normalized to *Ppia* and *Rps18* in QSCs, ASCs, myoblasts (MBs) and 3 days-differentiated muscle cells or myotubes (MTs) (n = 6 (3 males and 3 females for each genotype)). **d**) Representative immunofluorescence staining of PAX7 (red) labelling quiescent satellite cells on an isolated myofiber (T0 - Isolated myofiber), and primary myoblasts (T0 - Primary myoblasts) and reserve cells appearing along the myotubes (T72 - Myotubes). Myosin heavy chain (MyHC, green, asterisks) labels differentiated muscle cells or myotubes (MTs). **e**) Representative immunostaining of gamma-TUBULIN (γTUB, red) and acetylated alpha-TUBULIN (acαTUB, green) showing the basal body and primary cilium on a PAX7^+^ quiescent satellite cell (purple, T0 - QSC). γTUB (red) and ARL13B (green) label respectively the basal body and primary cilium of both PAX7^+^ myoblast (purple, T - MB) and reserve cell (purple, T72 - RC). Nuclei (blue) are labelled with DAPI. Scale bars, 10μm; Error bars, SEM; \**p* < 0.05.

**Supplementary Figure 2.**
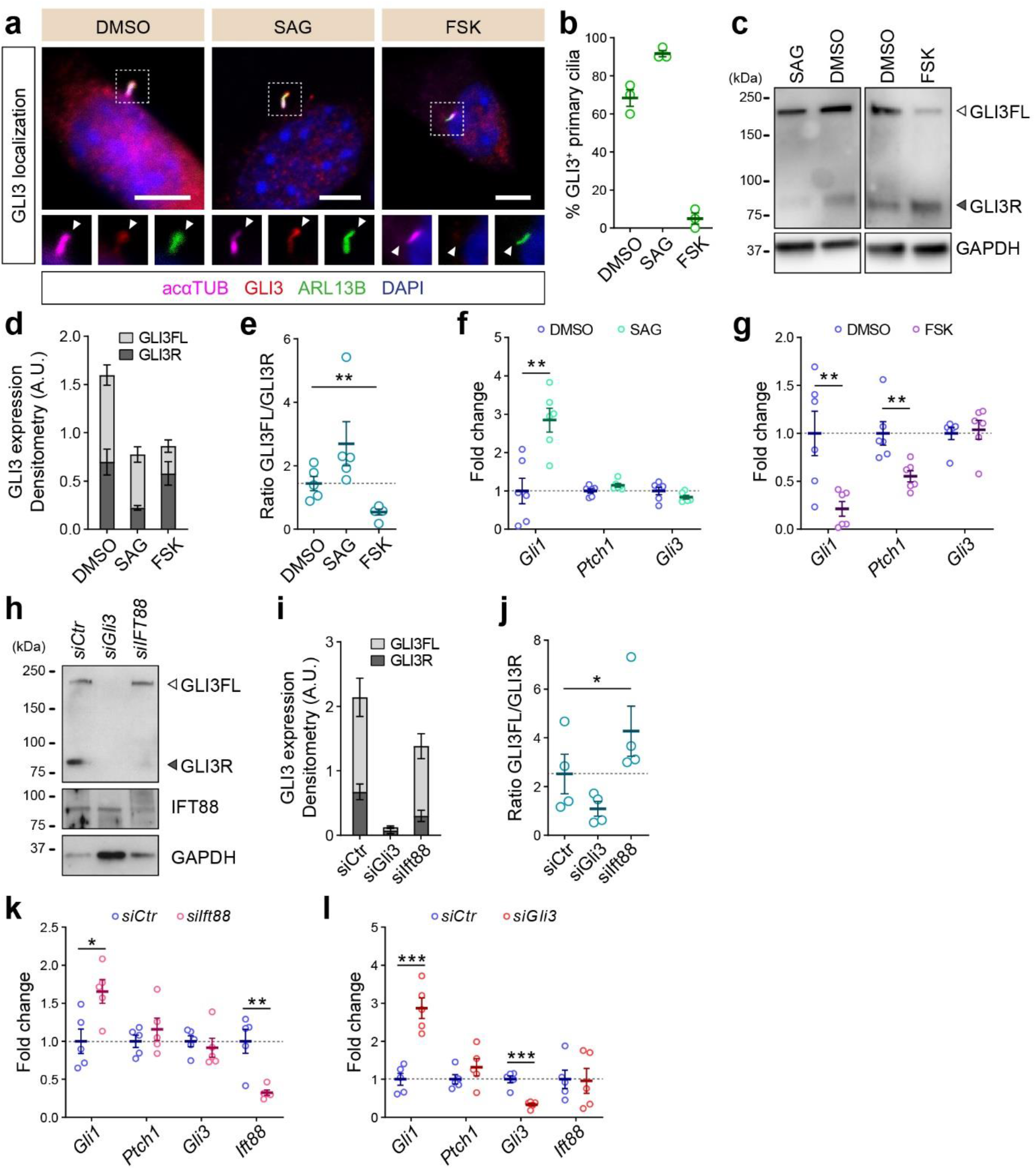
GLI3 processing and subcellular localization relies on the primary cilium and regulates Hedgehog signaling target genes. **a**) Representative immunofluorescence pictures showing GLI3 (red) localization in primary cilia stained with both acαTUB (purple) and ARL13B (green) upon SAG (Smoothened agonist) or FSK (forskolin) treatment. DMSO (vehicle) is used as a control. Scale bars, 5μm. **b**) Proportion of DMSO, SAG or FSK treated-myoblasts that exhibit GLI3 staining in the primary cilium (n = 3). **c**) Immunoblot analysis of GLI3 full-length (GLI3FL) and repressor (GLI3R) in DMSO, SAG or FSK treated-myoblasts. GAPDH is used as a loading control. **d**) Densitometric analysis of the level of GLI3FL (light gray) and GLI3R (dark gray) relative to GAPDH signals of 5 biological replicates. **e**) Ratio of GLI3FL/GLI3R relative to GAPDH (n = 5). **f**) Expression levels of *Gli1* and *Ptch1*, two Hh target genes, normalized to *Ppia* and *Gapdh*, upon SAG treatment, serving as a metric of canonical Hh pathway activation (n = 6). **g**) Expression levels of *Gli1* and *Ptch1*, two Hh target genes, normalized to *Ppia* and *Gapdh*, upon FSK treatment, showing the inhibition of canonical Hh signaling (n = 6). **h**) Immunblot analysis of GLI3 and IFT88 confirming the efficiency of *Gli3* and *Ift88* knockdown in primary myoblasts. **i**) Densitometric analysis of the level of GLI3FL (light gray) and GLI3R (dark gray) relative to GAPDH signals of 4 biological replicates. **j**) Ratio of GLI3FL/GLI3R relative to GAPDH (n = 4). **k**) Expression of *Gli1, Ptch1, Gli3* and *Ift88* normalized to *Ppia* and *Gapdh* in primary myoblasts, 48h after treatment with siRNA for *Gli3 (siGli3)* or with a non-target control *(siCtr)* (n = 5). **l**) Expression of *Gli1, Ptch1, Gli3* and *Ift88* normalized to *Ppia* and *Gapdh* in primary myoblasts, 48h after treatment with siRNA for *Ift88 (siIft88)* or with a non-target control *(siCtr)* (n = 5). Scale bars, 10μm; Error bars, SEM; \**p* < 0.5, \*\**p* < 0.01, \*\*\**p* < 0.001.

**Supplementary Figure 3.**
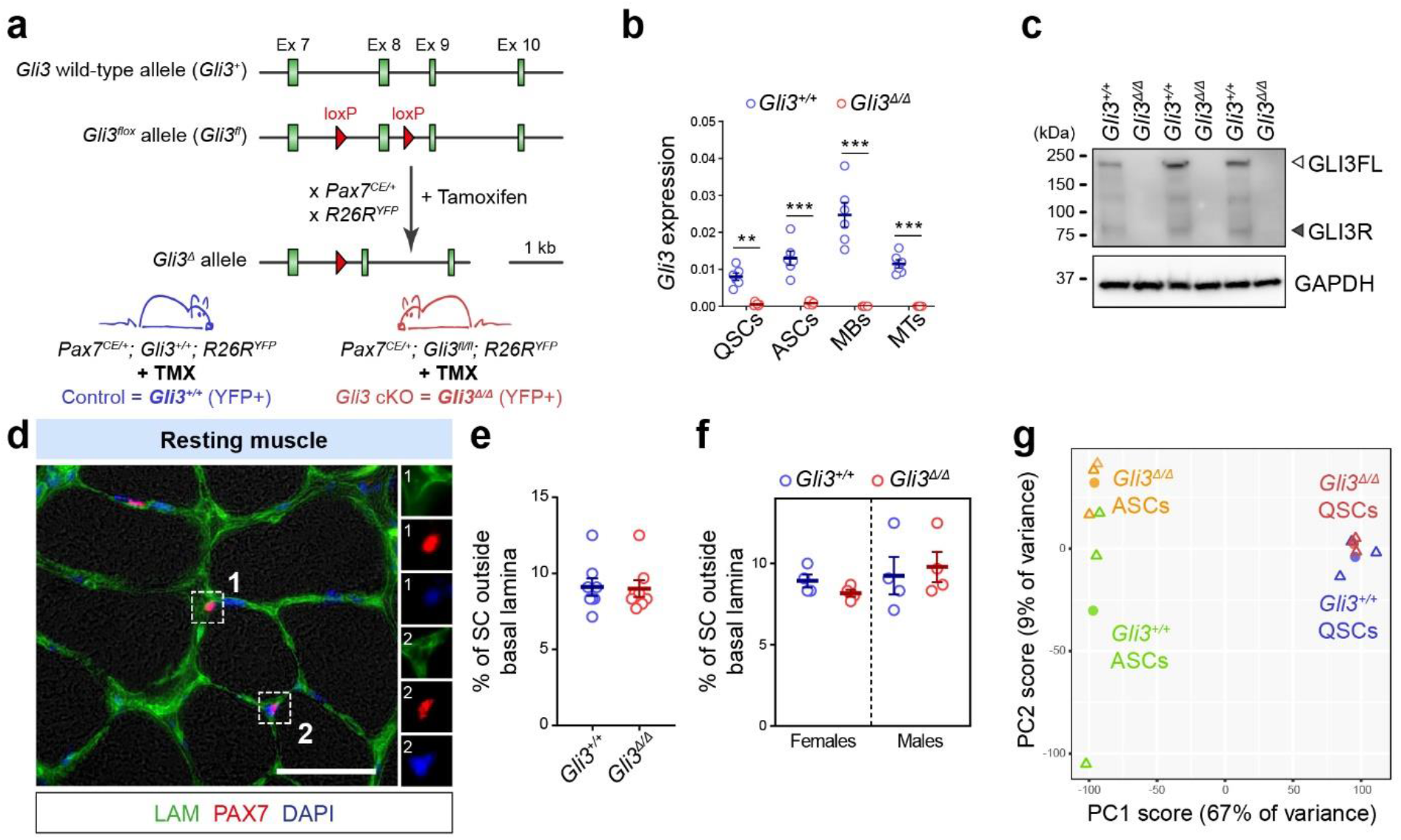
Resting muscle phenotype of the Pax7^CE/+^;Gli3^fl/fl^;R26R^YFP^ mice. **a**) The exon 8 of the *Gli3* floxed allele *(Gli3^fl^)* is flanked by two loxP sites, allowing for its tamoxifen (TMX)- inducible recombination by the CreER recombinase. The CreER inserted downstream the *Pax7* stop codon *(Pax7^CE/+^)* allows endogenous PAX7 expression while permitting specific ablation of *Gli3* in satellite cells upon TMX treatment. In addition, the *R26R^YFP^* allele was added allowing for tracing the satellite cell (SC) population. TMX-treated *Pax7^CE/+^; Gli3^+/+^; R26R^YFP^* mice are used as control *(Gli3^+/+^)* while TMX-treated *Pax7^CE/+^; Gli3^fl/fl^; R26R^YFP^* mice where SCs will be conditionally knocked-out for *Gli3* are referred as *Gli3*^Δ/Δ^. **b**) Expression analysis by RT-qPCR of *Gli3* normalized to *Ppia* and *Rps18* in quiescent (QSCs) and activated (ASCs) satellite cells, primary myoblasts (MBs), and 3 days differentiated myotubes (MTs) (n = 6). **c**) Western blotting for GLI3 confirming the knockout efficiency in primary myoblasts. GAPDH is used as a loading control (n = 3 biological samples). **d**) Immunofluorescence for PAX7 (red) and LAMININ (LAM, green) showing 1) a satellite cell within its niche and 2) a satellite cell surrounded by the basal lamina outside its niche. **e**) Proportion of satellite cells found outside the basal lamina in *Gli3*^+/+^ and *Gli3*^Δ/Δ^ resting TA muscles (n = 6, 3 males and 3 females for each genotype). **f**) Proportion of SCs found outside the basal lamina in *Gli3*^+/+^ and *Gli3*^Δ/Δ^ males and females (n = 3). **g**) Principal component analysis (PCA) of global transcriptomes of *Gli3*^+/+^ and *Gli3*^Δ/Δ^ quiescent satellite cells (QSCs) and activated satellite cells (ASCs). Each triangle represents a biological replicate. Each dot represents the mean of 3 biological samples (n =3 males for each condition and genotype).

**Supplementary Figure 4.**
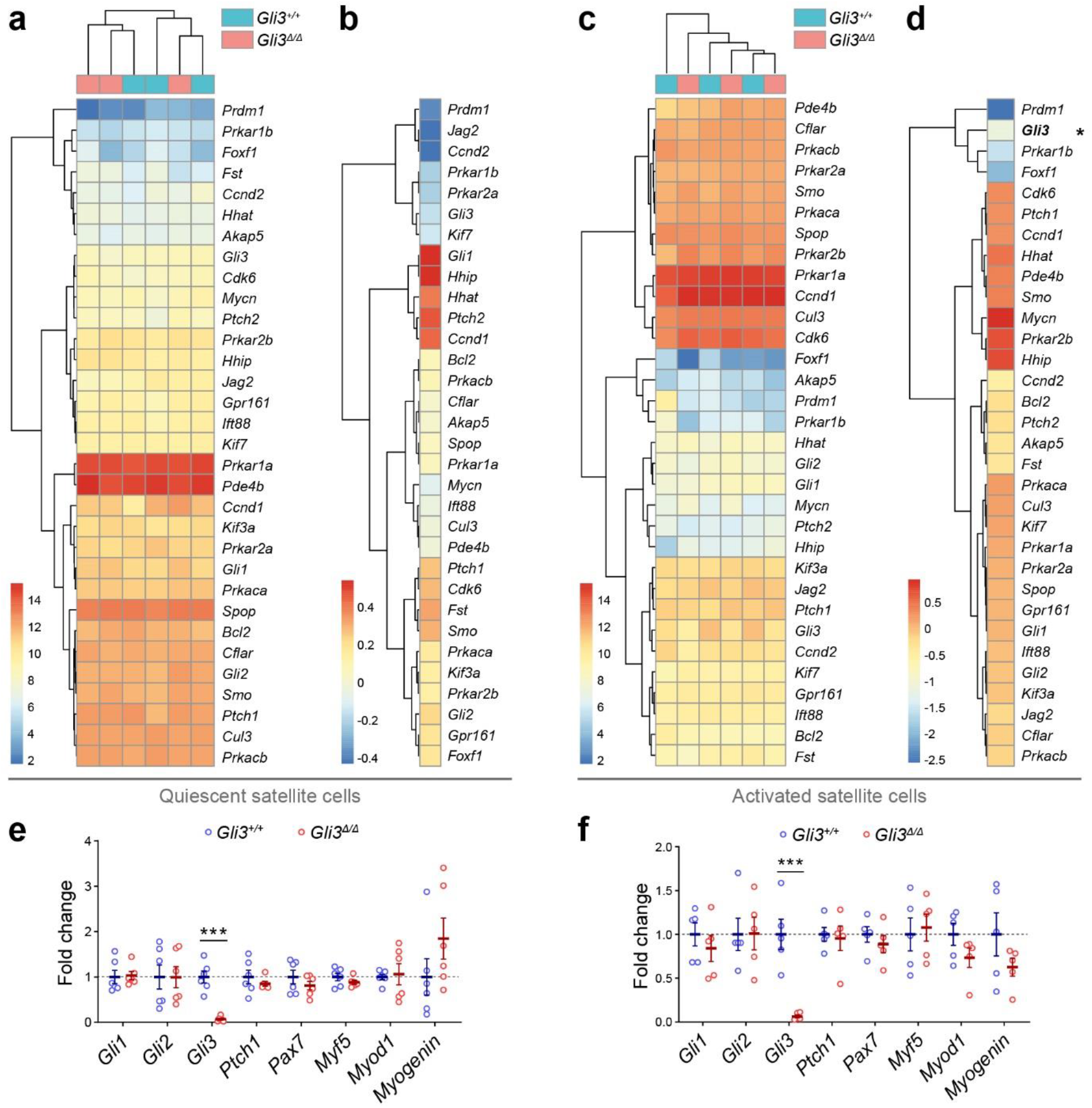
Genes of canonical Hedgehog signaling does not vary upon Gli3 deletion in QSCs and ASCs. **a**) Heatmap from normalized and log2 transformed expression matrix of components and target genes of canonical Hedgehog signaling in *Gli3*^Δ/Δ^ and *Gli3*^+/+^ QSCs (n = 3 males). **b**) Heatmap showing fold change values of canonical Hedgehog signaling genes in *Gli3*^Δ/Δ^ compared to *Gli3*^+/+^ QSCs. **c**) Heatmap from normalized and log2 transformed expression matrix of components and target genes of canonical Hedgehog signaling in *Gli3*^Δ/Δ^ and *Gli3*^+/+^ ASCs (n = 3 males). **d**) Heatmap showing fold change values of canonical Hedgehog signaling genes in *Gli3*^Δ/Δ^ compared to *Gli3*^+/+^ ASCs. Only *Gli3* (in bold, with a star) is significantly down-regulated with a padj < 0.05. **e**) RT-qPCR analysis of *Gli1-3*, *Ptch1*, *Pax7*, *Myf5*, *Myod1* and *myogenin* normalized to *Ppia* and *Gapdh* in *Gli3*^Δ/Δ^ and *Gli3*^+/+^ QSCs (n = 5). **f**) RT-qPCR analysis of *Gli1-3*, *Ptch1*, *Pax7*, *Myf5*, *Myod1* and *myogenin* normalized to *Ppia* and *Gapdh* in *Gli3*^Δ/Δ^ and *Gli3*^+/+^ ASCs (n = 5).

**Supplementary Figure 5.**
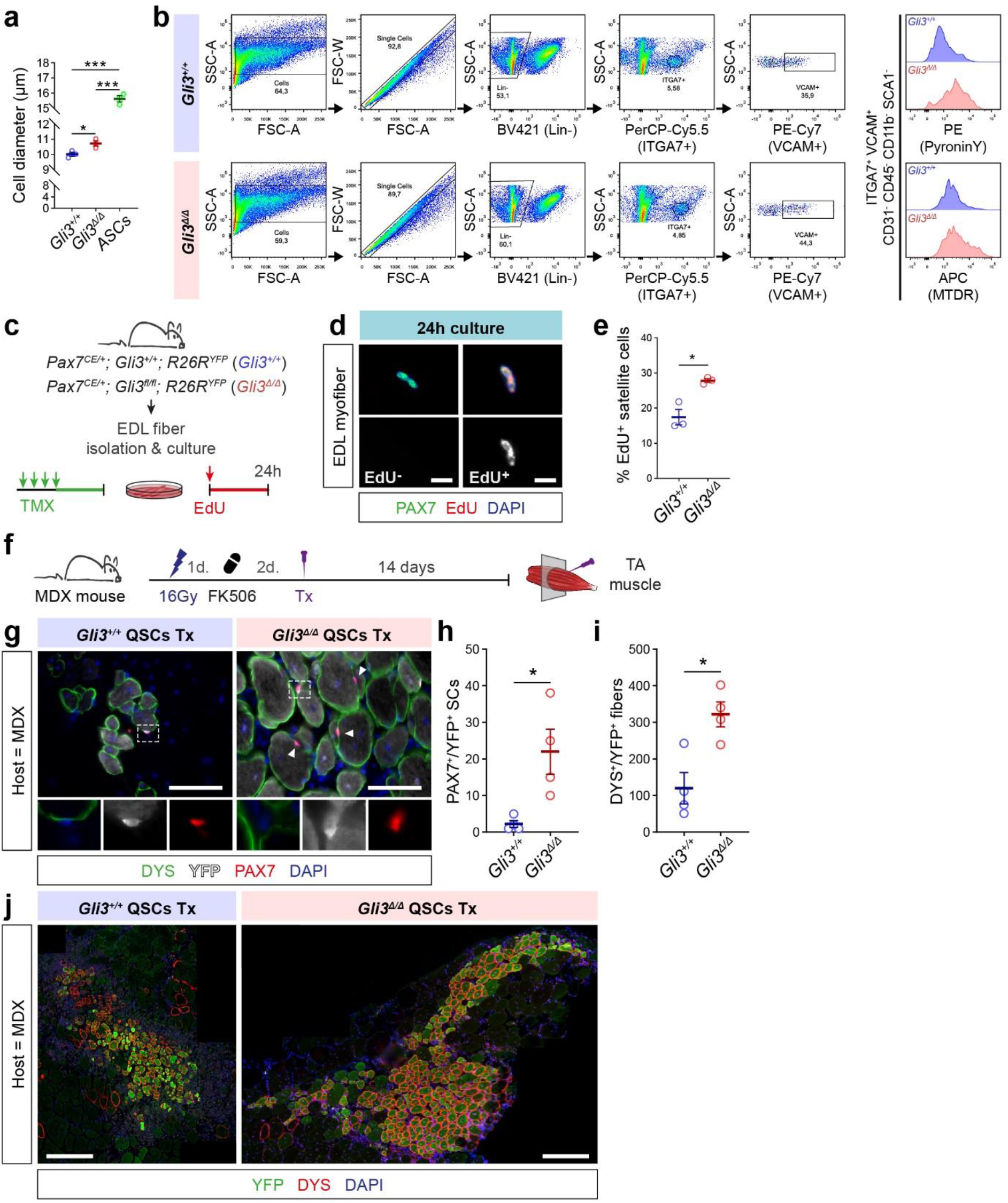
Gli3^Δ/Δ^ satellite cells exhibit G_Alert_ features, enter the cell cycle faster than Gli3^+/+^ satellite cells and engraft better. **a**) Mean cell diameter of *Gli3*^+/+^ and *Gli3*^Δ/Δ^ quiescent satellite cells (QSCs). ASCs were isolated from both *Gli3*^+/+^ and *Gli3*^Δ/Δ^ mice as they do not exhibit differences (n = 3 males). **b**) Flow cytometry gating strategy used to analyze PyroninY and MitoTracker Deep Red (MTDR) staining in QSCs from *Gli3*^+/+^ and *Gli3*^Δ/Δ^ mice. **c**) Experimental procedure followed to analyze the cell cycle entry of satellite cells from freshly isolated myofibers from *Gli3*^+/+^ and *Gli3*^Δ/Δ^ mice. **d**) Representative immunofluorescence of satellite cells (PAX7, green) that have incorporated EdU (EdU^+^, red) or not (EdU^-^) on 24h-cultured myofibers. Nuclei are stained with DAPI (blue). **e**) Proportions of EdU^+^ satellite cells 40h after isolation from *Gli3*^+/+^ and *Gli3*^Δ/Δ^ mice (n = 3 males). Scale bars, 10μm. **f**) 10,000 freshly isolated quiescent satellite cells (QSCs) from *Gli3*^+/+^ and *Gli3*^Δ/Δ^ mice were transplanted into TA muscles of 16Gy-irradiated and FK506-immunocompromised MDX mice. **g**) Immunofluorescence picture of transverse sections of MDX host TA muscle 14 days post-engraftment. Both satellite cells (PAX7, red) and myofibers (DYS, green) from the donors express YFP (grey). DAPI stains the nuclei. Scale bars, 50μm. **h**) Number of PAX7^+^/YFP^+^ satellite cells per transverse sections of MDX transplanted muscles. **i**) Number of DYS^+^/YFP^+^ myofibers per transverse sections of MDX transplanted muscles. **j**) Representative immunofluorescence picture showing the size of the engrafted area of MDX muscle 14 days following the transplantation of 10^4^*Gli3*^+/+^ or *Gli3*^Δ/Δ^ quiescent satellite cells (QSCs). Donor myofibers express both DYS (red) and YFP (green). DAPI stains the nuclei. Scale bars, 500μm. Unless indicated otherwise, n = 3 *Gli3*^+/+^ males and ≥ 3 *Gli3*^Δ/Δ^ males; Error bars, SEM; \**p* < 0.05; \*\*\**p* < 0.001.

**Supplementary Figure 6.**
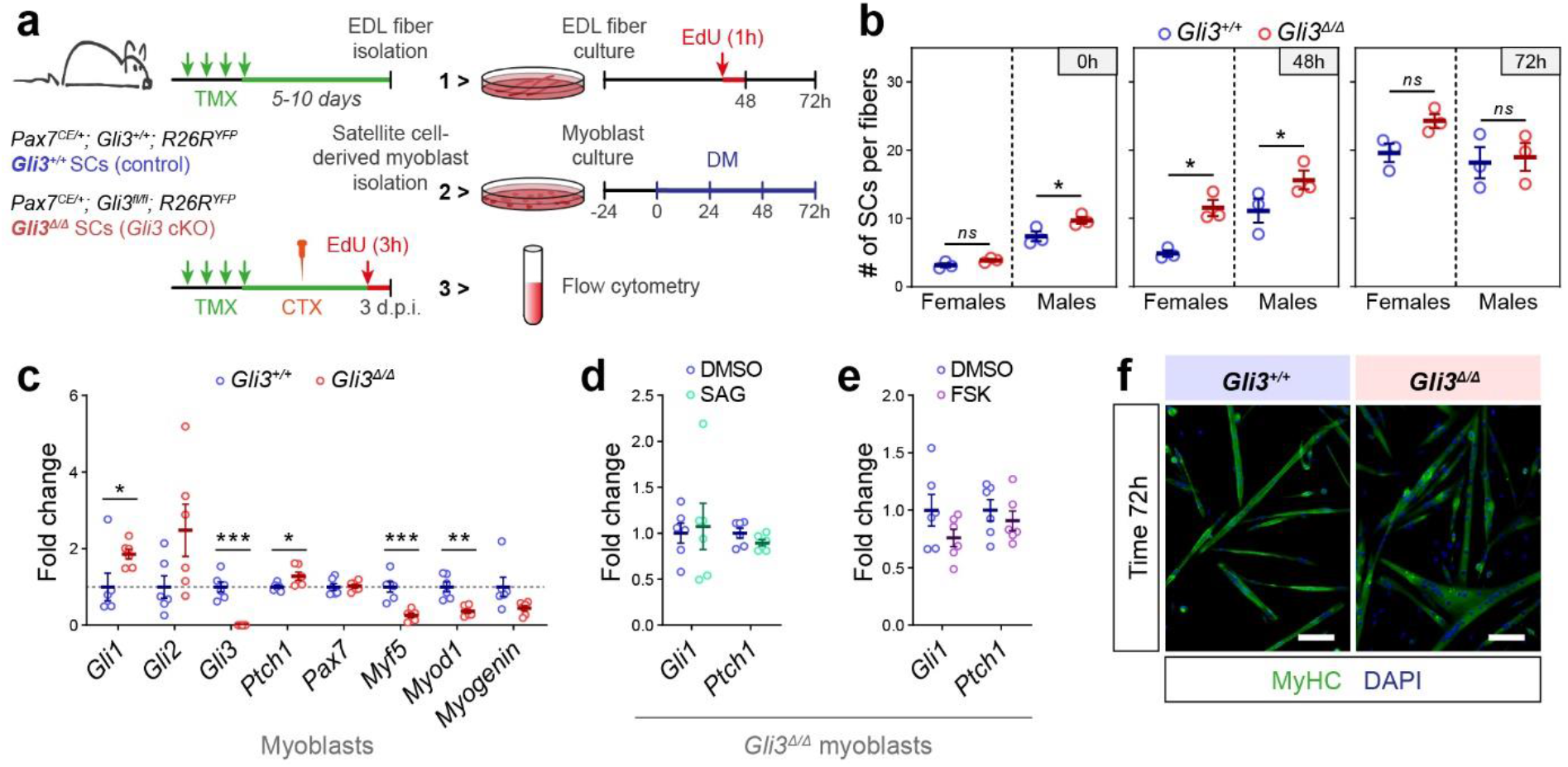
Phenotypic characterization of Gli3^+/+^ and Gli3^Δ/Δ^ activated satellite cells and primary myoblasts. **a**) Experimental design. **1>**Single EDL myofibers are isolated from tamoxifen-treated *Pax7^CE/+^;Gli3^+/+^;R26R^YFP^* (control, *Gli3^+/+^)* and *Pax7^CE/+^;Gli3^fl/fl^;R26R^YFP^ (Gli3* conditional knockout, *Gl3^Δ/Δ^)* mice and cultured for 48h and 72h to follow satellite cell proliferation and differentiation. **2>** Satellite cell-derived myoblasts were cultured in proliferating conditions and differentiated for 72h in differentiation medium (DM). **3>** TA and *gastrocnemius*(GA) muscle injury was induced by cardiotoxin (CTX) injection. 3 days post-injury (3 d.p.i.), activated satellite cells (ASCs) were FACS-isolated and processed for RNA extraction (RNA-sequencing and qPCR). **b**) Quantification of the number of satellite cells per myofiber immediately after isolation (0h), or after 48h and 72h of culture (n = 3 males and 3 females for each genotype). **c**) RT-qPCR analysis of *Gli1-3*, *Ptch1*, *Pax7*, *Myf5*, *Myod1* and *myogenin* normalized to *Ppia* and *Gapdh* in *Gli3*^Δ/Δ^ and *Gli3*^+/+^ proliferating primary myoblasts (n = 6). **d**) Expression of *Gli1* and *Ptch1* normalized to *Ppia* and *Gapdh*, in *Gli3*^Δ/Δ^ myoblasts upon SAG treatment (n = 6). **e**) Expression of *Gli1* and *Ptch1* normalized to *Ppia* and *Gapdh*, in *Gli3*^Δ/Δ^ myoblasts upon FSK treatment (n = 6). **f**) Immunostaining of myosin heavy chains (MyHC, green) of 72h-differentiated *Gli3*^Δ/Δ^ and *Gli3*^+/+^ myoblasts. Nuclei are counterstained with DAPI (blue). Scale bar, 100μm. Error bars, SEM; \**p* < 0.05; \*\**p* < 0.01; \*\*\**p* < 0.001.

**Supplementary Figure 7.**
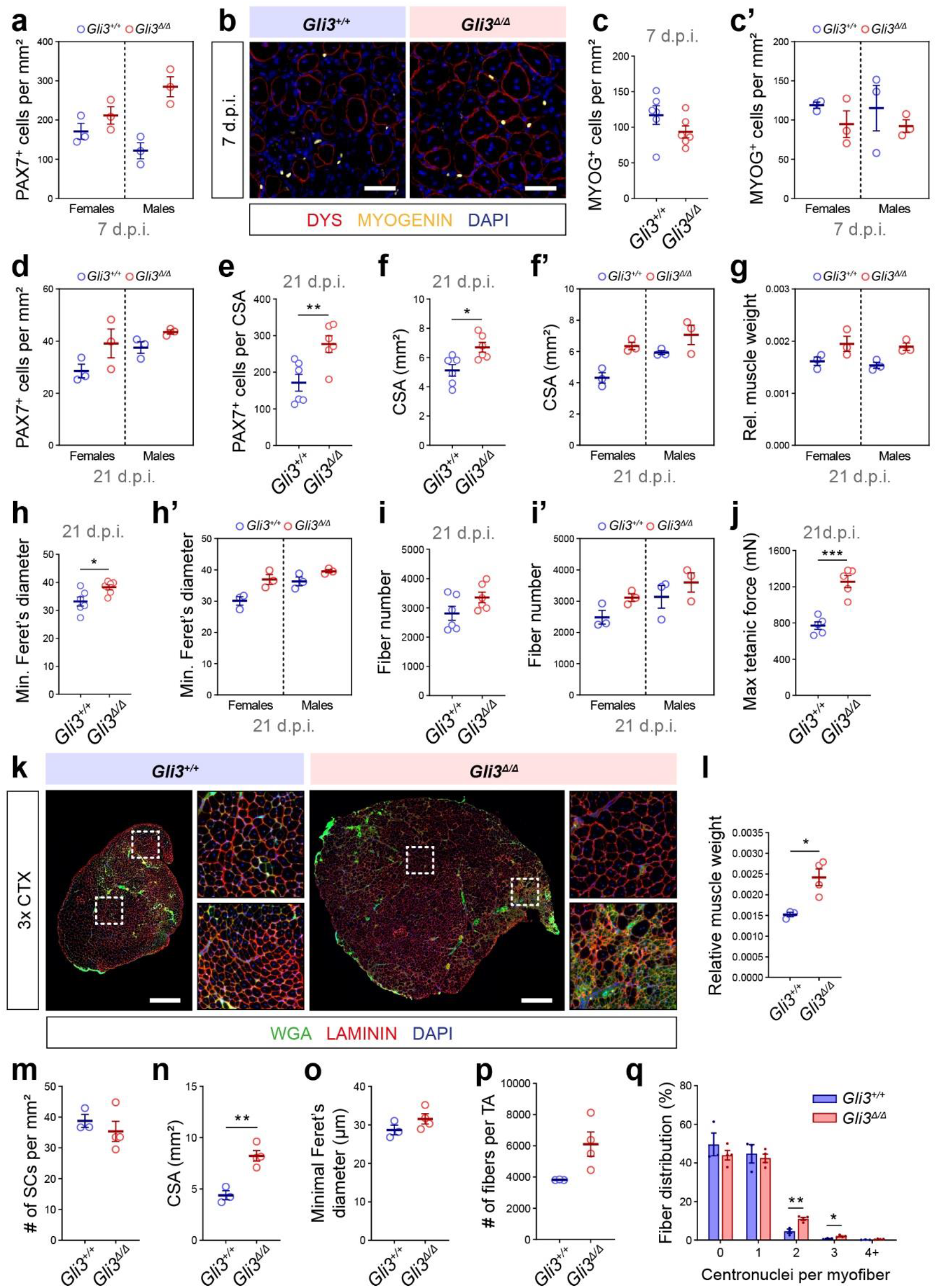
Phenotype of Gli3^+/+^ and Gli3^Δ/Δ^ muscle following a single and triple cardiotoxin-injuries. **a**) Quantification of PAX7^+^ cells per mm^2^ TA section of *Gli3*^+/+^ and *Gli3*^Δ/Δ^ female and male mice at 7 d.p.i. **b**) Immunostaining of MYOGENIN (yellow) at 7 d.p.i. showing the differentiated muscle cells and DYSTROPHIN (DYS, red) delineating the regenerating myofibers. DAPI stains the nuclei (blue). **c**) Quantification of MYOG^+^ cells per mm^2^ TA section of all *Gli3*^+/+^ and *Gli3*^Δ/Δ^ mice at 7 d.p.i. and **c’**) in females and males. **d**) Quantification of PAX7^+^ cells per mm^2^ TA section *Gli3*^+/+^ and *Gli3*^Δ/Δ^ female and male mice at 21 d.p.i. **e**) Quantification of PAX7^+^ cells per TA cross-sectional area (CSA). **f**) CSA of TA section of all *Gli3*^+/+^ and *Gli3*^Δ/Δ^ mice at 21 d.p.i. and **f’**) in females and males. **g**) Relative muscle weight of *Gli3*^+/+^ and *Gli3*^Δ/Δ^ female and male mice at 21 d.p.i. **h**) Minimal Feret’s diameter of all *Gli3*^+/+^ and *Gli3*^Δ/Δ^ mice at 21 d.p.i. and **h’**) in males and females. **i**) Number of myofibers per TA section of all *Gli3*^+/+^ and *Gli3*^Δ/Δ^ mice at 21 d.p.i. and **i’**) in males and females. **j**) Maximum tetanic force of TA muscles of *Gli3*^+/+^ and *Gli3*^Δ/Δ^ mice at 21 d.p.i. (n = 5 males). Unless indicated otherwise, n = 6 (3 males and 3 females for each genotype); Scale bars, 50μm; Error bars, SEM; *\*p* < 0.05; *\*\*p* < 0.01; *\*\*\*p* < 0.001. **k**) Representative immunofluorescence picture of *Gli3*^+/+^ or *Gli3*^Δ/Δ^ regenerated muscle following a triple injury (3x CTX). LAMININ (red) and WGA (green) delineate the myofibers. DAPI stains the nuclei. **l**) TA muscle weight normalized to total body weight of *Gli3*^+/+^ and *Gli3*^Δ/Δ^ mice following a triple injury. **m**) Quantification of satellite cells per mm^2^ 3x-injured TA muscle section of *Gli3*^+/+^ and *Gli3*^Δ/Δ^ mice. **n**) Crosssectional area (CSA) of 3x-injured TA muscle section of *Gli3*^+/+^ and *Gli3*^Δ/Δ^ mice. **o**) Minimal Feret’s diameter of regenerated myofibers of 3x-injured *Gli3*^+/+^ and *Gli3*^Δ/Δ^ mice. **p**) Number of myofibers per TA section of *Gli3*^+/+^ and *Gli3*^Δ/Δ^ mice after a triple injury. **q**) Distribution of regenerated myofibers according to their number of centrally located nuclei (centronuclei) of 3x-injured *Gli3*^+/+^ and *Gli3*^Δ/Δ^ mice. For the triple injuries (3x CTX), n = 3 *Gli3*^+/+^ males and 4 *Gli3*^Δ/Δ^ males; Scale bars, 500μm; Error bars, SEM; \**p* < 0.05; \*\**p* < 0.01.

